# Protein folding stress shapes microglial phenotype in progressive supranuclear palsy

**DOI:** 10.64898/2026.01.26.700084

**Authors:** Satoshi Tanikawa, Takeru Kuwabara, Shinya Tanaka, Gabor G. Kovacs

## Abstract

Microglia play a pivotal role in neurodegeneration, yet their response to tau pathology remains incompletely understood. Through single-nucleus RNA sequencing of Aβ plaque–free frontal cortex from 4R tauopathy progressive supranuclear palsy (PSP) brains, we reveal a distinct transcriptional reprogramming of homeostatic microglia. PSP microglia exhibit pronounced protein-folding and ER-stress signatures, elevated homeostatic and MHC class II gene expression, and attenuated cytoskeletal, motility, and interferon-related programs, accompanied by reduced inferred intercellular communication. Neuropathological analysis corroborates a shift toward highly ramified morphologies and increased MHC class II positivity in PSP cortex. In HMC3 cells, pharmacological induction of protein-folding stress recapitulates this phenotype, driving homeostatic gene upregulation and morphological transition to process-bearing forms. We define this protein-folding–stress–associated microglial (PSAM) state as a PSP-linked phenotype mechanistically distinct from disease-associated microglia (DAM), highlighting ER stress as a key driver of microglial remodeling in tauopathies.

## INTRODUCTION

Microglia, the resident immune cells of the brain, are essential for maintaining neuronal health and responding to injury. Recent advances in single-cell and single-nucleus transcriptomics have revealed that microglia are highly plastic, adopting distinct transcriptional states in response to specific microenvironmental cues^1,2^. In neurodegenerative research, current concepts have been strongly shaped by work in amyloid-driven Alzheimer’s disease (AD) experimental models. Studies in 5xFAD and related mice identified “disease-associated microglia” (DAM)^3^—also termed MGnD^4^ or ARM^5^—characterized by downregulation of homeostatic genes (for example, *P2RY12*, *CX3CR1*) and upregulation of genes involved in phagocytosis, lysosomal function, and lipid metabolism^3,6^. These cells preferentially cluster around extracellular amyloid plaques and are thought to help contain and clear protein aggregates^3,6^. Although human–mouse differences are evident^7^, these plaque-associated signatures have often been regarded as a prototypical microglial response to neurodegeneration^8^. It remains unclear, however, whether this plaque-centric DAM framework applies to tau-driven disorders that lack extracellular filamentous Amyloid beta filaments. Indeed, human tissue-based studies indicate distinct microglial states linked to amyloid-β (AD1) versus tau pathology (AD2), with AD1 resembling activated, phagocytic profiles from amyloid-based mouse models and AD2 defining a previously unrecognized state^19^.

Based on these we focused on progressive supranuclear palsy (PSP), an amyloid-independent 4R tauopathy characterized by the accumulation of abnormal tau in both neurons and glia^9,10^, providing an opportunity to dissect microglial responses to tau pathology in a tau-predominant, plaque-negative setting. Here we profiled microglia in amyloid plaque–negative PSP cortex using single-nucleus RNA sequencing, regulon analysis, histopathology, and *in vitro* assays. We identify a PSP-enriched state with ER-stress signatures and preserved homeostatic features, termed protein-folding–stress–associated microglia (PSAM), a program distinct from plaque-proximal DAM.

## RESULTS

### Quality Control in snRNA-seq Analysis of Human Brain Samples

We performed snRNA-seq on frozen frontal lobe specimens from PSP (n=8) and non-diseased control (n=5) subjects (Figure 1A). In the brain, neurons have higher RNA content and detected gene counts per nucleus compared to glia^11^, while microglial nuclei have relatively low counts. Due to the significant impact of neuron-derived ambient RNA and doublets^12^, we implemented background removal using CellBender^13^, based on the number of RNAs and the ratio of mitochondrial RNAs, and doublet removal using DoubletFinder^14^ (Figure S1A). Subsequently, cell annotation was performed based on the human motor cortex reference of Azimuth^15^ (Figure S1B). This revealed that multiple cell types were mixed in various proportions within single Seurat clusters (Figure S1C and E). We designated clusters composed of 75% of a single cell type as high-confidence clusters and excluded the remaining mixed cell-type clusters as “undetermined” (Figure 1D).

**Figure 1.**
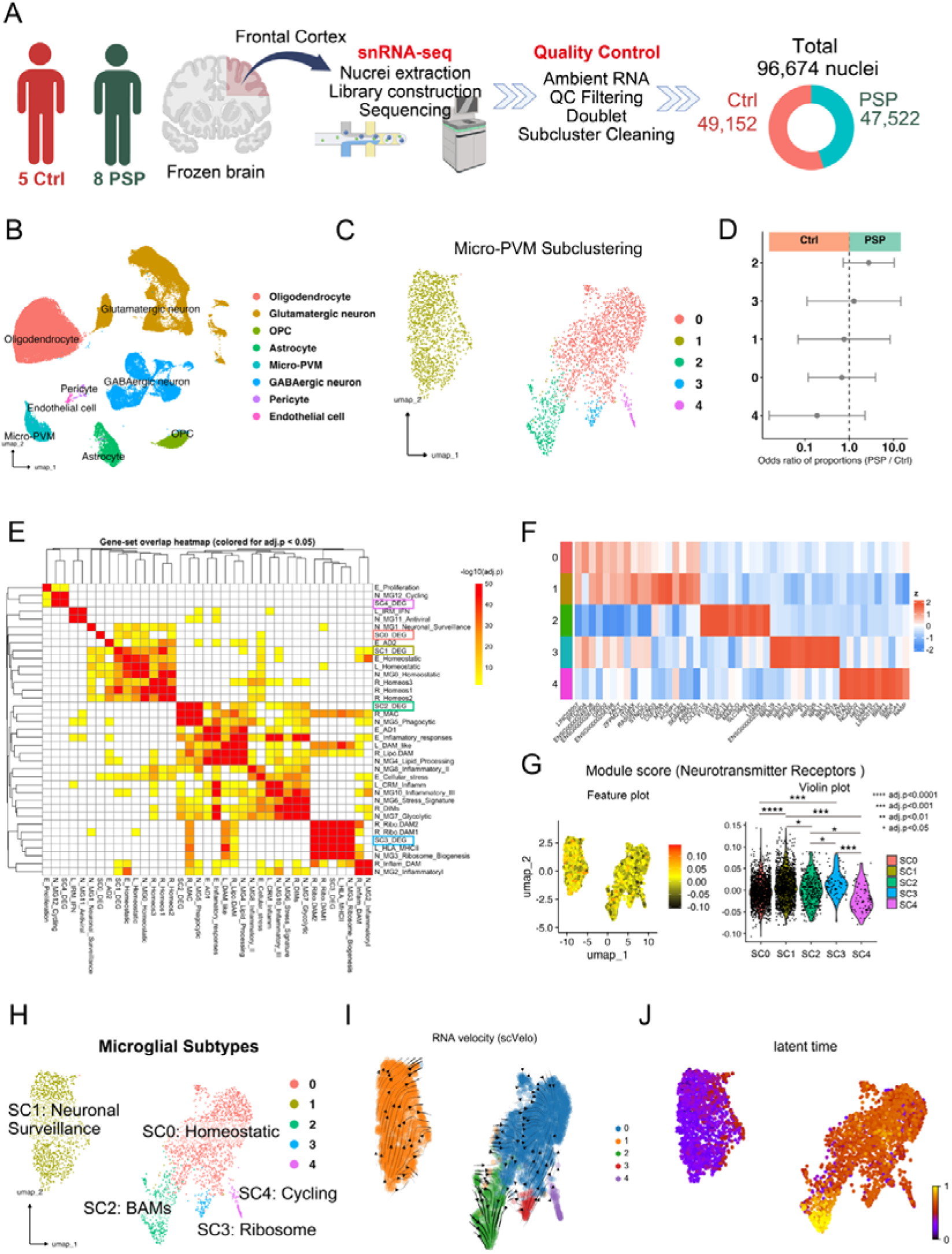
**Characteristics of microglial subclusters** (A) snRNA-seq flow. Created in BioRender. (B) UMAP plot of all cell types after quality control and integration. (C) UMAP visualization of subclustering within the microglia–perivascular macrophage (PVM) compartment. Each dot represents a single cell, and colors indicate five transcriptionally distinct subclusters (SC0–SC4). (D) Forest plot of cell proportions. Dots indicate the odds ratio of cell proportions (PSP / Ctrl) for each subcluster, and horizontal bars represent the 95% confidence intervals on a log_10_ scale; values >1 indicate relative enrichment in PSP, whereas values <1 indicate relative enrichment in controls. None of the subclusters showed a statistically significant difference in abundance between PSP and controls (adjusted p < 0.05). (E) Heatmap of gene-set overlap between previously reported microglial gene sets and DEGs from each subcluster. The matrix includes 32 published microglial gene sets (defined by upregulated genes) together with subcluster-specific DEG sets (SC0–SC4); pairwise overlap was quantified using the Jaccard index and hierarchically clustered. Statistical significance of overlap was assessed by one-sided Fisher’s exact tests with Benjamini–Hochberg correction, and only significant overlaps (adjusted p < 0.05) are colored according to –log_10_(adjusted p). (F) Heatmap of up to the top 10 DEGs per subcluster identified by pseudobulk analysis using glmQLFTest. Expression values are scaled by Z-score across samples, and genes that overlapped between subclusters are shown only once so that each column represents a unique gene. (G) UMAP feature plot and violin plots showing per-cell module scores for the neurotransmitter receptor gene set (**** adj. p < 0.0001, *** adj. p < 0.001, * adj. p < 0.05). (H) UMAP visualization of microglial subclusters with assigned subtype labels. (I) RNA velocity field for microglial subclusters inferred with scVelo. The UMAP embedding shows SC0–SC4 colored as indicated, with overlaid arrows representing the direction and magnitude of predicted transcriptional state transitions. (J) Latent time analysis inferred from RNA velocity. Each nucleus is colored according to its latent time (purple to yellow).

Next, we identified and removed ambient clusters following the procedure proposed by Caglayan et al.^12^(Figure S1A). We identified 178 ambient markers; as noted by Caglayan et al., a large portion consisted of neuron-derived ambient RNA, with 38% (65/178) being of neuronal origin (Data S2). Within the microglia subclusters, we observed subclusters highly expressing oligodendrocyte-derived markers (*MOG*, *MBP*, *MOBP*) and neuron-derived markers (*SYT1*, *CSMD1*, *KCNIP4*) (Figure S2A and 2B). Subclusters with a high expression of ambient RNAs compared to other subclusters were identified as ambient clusters and removed (Figure S2C). Using the same procedure, we removed ambient clusters from all cell types except for Glutamatergic and GABAergic neurons (Figure S2D). Finally, 96,674 high-quality nuclei (PSP=47,522, Ctrl=49,152) were integrated and used for downstream analysis (Figure 1B).

### Five microglial subclusters across PSP and control frontal cortex

We analyzed a total of 3,499 microglial nuclei after quality control (PSP, 2,697; Ctrl, 802). Five subclusters were identified by subclustering (Figure 1C). Differential abundance analysis showed no subcluster (SC) with a significantly increased or decreased proportion in PSP compared with controls (Figure 1D). Point estimates suggested a mild enrichment of SC2 in PSP and of SC4 in controls, but these trends did not reach statistical significance. To characterize each SC, we compiled 32 gene sets from four published human brain snRNA-seq studies^16–19^ (Data S5). We first performed pseudobulk differential expression analysis to obtain subcluster-specific differentially expressed genes (DEGs) and assessed overlap with reported gene sets. Set similarity was quantified by the Jaccard index, and hierarchical clustering was based on Jaccard distance (Data S5). Significance of overlap was evaluated by Fisher’s exact test with Benjamini–Hochberg correction (Data S5). Clustering of Jaccard overlap between published gene sets organized them into biologically coherent modules, including a cluster of homeostatic signatures and a distinct cluster of DAM-related signatures, consistent with their original labels (Figure 1E).

SC0 and SC1 both showed high expression of canonical homeostatic microglial markers (*P2RY12* and *SALL1*) and clustered within the homeostatic gene׈set module. SC0 showed particularly strong overlap with N_MG0_Homeostatic (Na Sun) and E_Homeostatic (Emma Gerits), whereas SC1 overlapped significantly with all homeostatic gene sets and additionally with N_MG1_Neuronal_Surveillance, E_Cellular_stress and other inflammatory signatures. SC0 and SC1 also shared two of the top ten upregulated DEGs identified by a conservative procedure (*ENSG00000249738* and *XACT*), and their overall expression profiles were highly similar (Figure 1F). Consistently, modulelscore analyses showed higher N_MG1_Neuronal_Surveillance scores in SC1 than in SC0, together with lower AD1/AD2 scores (Figure S3 and Data S3). The “Neurotransmitter receptor” module, a key component of N_MG1_Neuronal_Surveillance, was highest in SC1 and significantly higher than in SC0, SC2 and SC4 (Figure 1G and Data S5). Thus, while SC0 and SC1 both represent homeostatic׈like microglia, SC1 additionally displays pronounced neuronal׈surveillance features, and we therefore annotated SC0 as Homeostatic and SC1 as Neuronal surveillance.

The DEGs of SC2 included several border-associated macrophage (BAM)–related genes such as *CD163*, *MRC1* and *LYVE1* (Data S5). Fisher’s exact test revealed strong and significant overlap with macrophage-related modules R_MAC and N_MG5_Phagocytic (Figure 1E and Data S5), and module scores for macrophage-related gene sets were significantly higher in SC2 than in the other clusters (Figure S3 and Data S3). These findings indicate that SC2 has macrophage-like properties, and we therefore annotated it as SC2: BAMs. The DEGs of SC3 were dominated by ribosomal protein genes (*RPS* and *RPL* families), which comprised most of its DEGs (59/89). These genes showed strong overlap with modules enriched for ribosomal components, including L_HLA_MHCII, N_MG3_Ribosome_Biogenesis and R_Ribo.DAM1/2 (Figure 1E), and module scores for ribosome-related gene sets were significantly higher in SC3 than in the other clusters (Figure S3 and Data S3). We therefore annotated SC3 as Ribosome. In SC4, DEGs included DNA-replication–related genes such as *POLE*, *POLD3* and *POLA1/2*, which significantly overlapped with N_MG12_Cycling and E_Proliferation (Figure 1E).

Consistently, module scores for E_Proliferation and M_Cell cycle were significantly higher in SC4 than in the other clusters (Figure S3 and Data S3), and we therefore annotated SC4 as Cycling. Taken together, these analyses defined five transcriptionally distinct microglial subtypes: SC0 (Homeostatic), SC1 (Neuronal surveillance), SC2 (BAMs), SC3 (Ribosome) and SC4 (Cycling), which occupied partially segregated regions on the Uniform Manifold Approximation and Projection (UMAP) embedding (Figure 1D).

RNA velocity analysis using scVelo^20^ suggested that transcriptional flows originated in SC1 and were directed towards both SC0 and SC2 (Figure 1I). Latent time analysis was consistent with this pattern, placing SC1 at early and SC2 at late latent times (Figure 1J). In addition to the terminal focus in SC2, the velocity field also showed a second convergence point within SC0, receiving vectors from SC1, SC3 and SC4, suggesting the presence of two major trajectories terminating in SC0 (Figure 1I).

### Homeostatic microglia (SC0) show the largest PSP–control divergence

Cellular proportions of microglial subclusters did not differ significantly between PSP and controls (Figure 1D). We therefore asked whether PSP and control nuclei might instead occupy different transcriptional states within the same subclusters. For each subcluster, we computed the Euclidean distance in UMAP space between the centroids of PSP and control nuclei. This analysis revealed the largest PSP–control separation for SC0 (Homeostatic; centroid distance = 0.80), with more modest shifts in SC2, SC1 and SC4 and minimal separation in SC3 (Figure 2A and 2B). Within SC0, the PSP centroid was located closer to the apparent convergence point of the velocity field, whereas the control centroid was displaced toward the SC2_BAMs region (Figure 1I).

**Figure 2.**
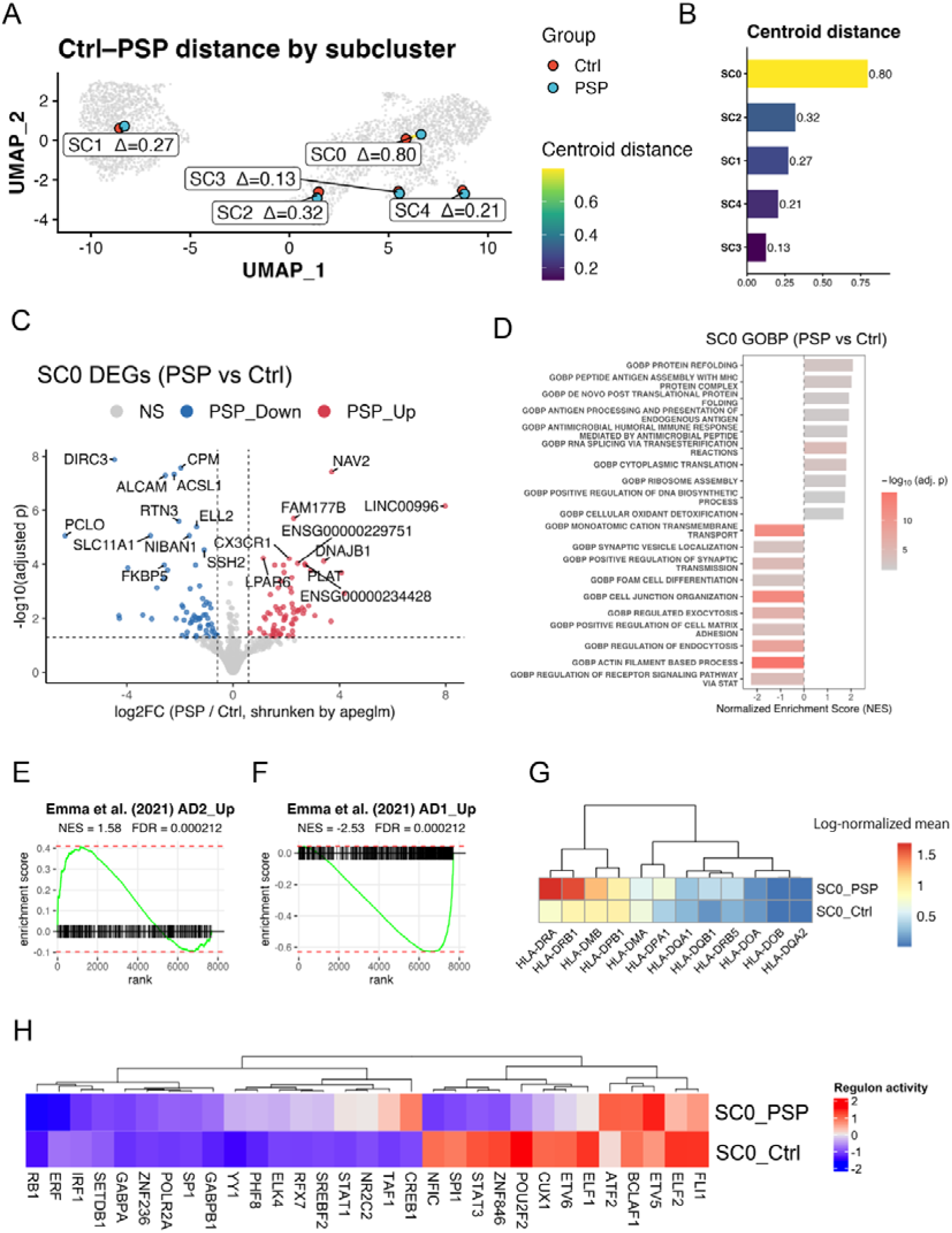
**PSP–control divergence in homeostatic microglia (SC0)** (A) Ctrl–PSP centroid distances across microglial subclusters. For each subcluster (SC0–SC4), the centroids of control and PSP cells were calculated in UMAP space, and the Euclidean distance between them (Δ) was used as a measure of transcriptional divergence. (B) The bar plot summarizing centroid distances. (C) Volcano plot of DEGs in SC0 (PSP vs control). Pseudobulk differential expression was performed with DESeq2 using apeglm shrinkage of log_2_ FC. Genes with adjusted p < 0.05 and absolute log_2_FC ≥ 0.25 are highlighted (red, upregulated in PSP; blue, downregulated). (D) Gene set enrichment analysis of the homeostatic subcluster SC0 (PSP vs control). Genes were ranked according to the pseudobulk DE results from DESeq2, and pathway enrichment was assessed with fgsea using the C5 Gene Ontology Biological Process (GOBP) collection. Bars show significantly enriched pathways (adjusted p < 0.05), with the normalized enrichment score (NES). (E and F) Gene set enrichment plots for the microglial signatures AD2_up (E) and AD1_up (F) from Emma et al. (2021), using the SC0_PSP-versus-Ctrl ranked gene list. (G) Heatmap of representative HLA/MHC class II–related genes in SC0. Rows show mean log-normalized expression in SC0_PSP and SC0_Ctrl; colors indicate expression level. (H) SCENIC-based transcription factor regulon activity in SC0. Heatmap shows z-scored activity for representative regulons in SC0_PSP and SC0_Ctrl, with red and blue indicating higher and lower activity, respectively.

Given this pronounced withinlcluster shift, we next split SC0 nuclei into PSP and control groups (SC0_PSP and SC0_Ctrl) and performed pseudobulk differential gene expression analysis. SC0_PSP showed 76 upregulated and 86 downregulated genes (Figure 2C and Data S6). Notably, upregulated genes included homeostatic microglial markers (*CX3CR1*, *GPR34* and *P2RY13*) and heat shock protein genes (*HSP90AA1*, *HSPB1*, *HSPE1* and *DNAJB1*). Downregulated genes included components of actin remodeling and cytoskeletal scaffolding (*DOCK3*, *SIPA1L1*, *ACTN1*, *FMN1*, *SSH2*, *SCIN*, *MYO1E*, *SH3PXD2B* and *NCK2*), adhesion molecules (*ALCAM* and *CADM1*), and receptor recycling/membrane trafficking (*TBC1D16*, *TBC1D8* and *DENND3*). In addition, interferon-pathway components (*IFI44*, *IFI44L*, *IFNAR2* and *IFNLR1*) were reduced. In gene set enrichment analysis of SC0_PSP versus SC0_Ctrl, protein-folding–related pathways were prominently enriched, with heat-shock protein genes represented in the leading edge. These included protein refolding, de novo post-translational protein folding (Gene Ontology biological process, GO:BP) and HSF1 activation (Reactome) (Figure 2D, Figure S4A, Data S4, and Data S6). Consistent with this, an ER-stress pathway, “ATF4 activates genes in response to endoplasmic reticulum stress” (Reactome), was also significantly enriched among PSP-upregulated genes (Figure S4A and Data S4). Pathways related to antigen processing and presentation were likewise detected, with HLA-DRA among the leading-edge genes, for example peptide antigen assembly with MHC protein complex and antigen processing and presentation of endogenous antigen (GO:BP). In contrast,

PSP-downregulated genes were dominated by pathways involved in the cytoskeleton, adhesion and motility, including actin filament–based processes and cell–cell adhesion (GO:BP) and the CDC42 and RHO GTPase cycles (Reactome). Additional pathways associated with membrane trafficking and immune signaling were suppressed, such as interferon-mediated signaling, endocytosis and regulated exocytosis (GO:BP). In a complementary Metascape^21^ analysis of SC0_PSP DEGs, the Reactome term “GPCR ligand binding”, which includes several homeostatic microglial receptors, was significantly enriched, and Gene Ontology terms related to “response to unfolded protein” and “regulation of chemotaxis” ranked among the top pathways (Figure S4B and Data S4). Consistent with the GSEA results, pathways associated with the actin cytoskeleton, cell junction assembly and interferon-stimulated responses showed negative enrichment in SC0_PSP (Figure S4B and Data S4). Taken together with the GSEA findings, these results indicate that SC0_PSP exhibits relatively stronger signatures related to GPCR/homeostatic signaling and protein folding/ER stress, whereas pathways involved in cytoskeletal regulation, interferon signaling, and endocytosis/exocytosis are attenuated.

Using a preranked gene list for SC0_PSP versus SC0_Ctrl, we performed GSEA against the plaque-associated (E_AD1) and tau-related microglial (E_AD2) signatures reported by Emma et al. E_AD2 was significantly positively enriched (NES = 1.58, FDR = 2.1 × 10ll;Figure 2F), whereas E_AD1 was significantly negatively enriched (NES = −2.53, FDR = 2.1 × 10ll; Figure 2G). These results indicate that SC0_PSP aligns more closely with a tau-pathology–related microglial program and, conversely, diverges from a plaque-proximal DAM-like program.

Pathways related to antigen processing and presentation also distinguished SC0_PSP from the homeostatic SC0_Ctrl state. Within SC0, HLA class II genes were consistently expressed at higher levels in PSP than in controls, with particularly marked increases in *HLA-DRA* and *HLA-DRB1* (Figure 2H). Notably, this upregulation of MHC class II occurred despite the preserved high expression of homeostatic microglial markers (*CX3CR1*, *GPR34* and *P2RY13*), indicating a hybrid state combining homeostatic and activated features. This pattern is concordant with the GSEA enrichment of antigen presentation pathways.

Gene regulatory network analysis was performed using single-cell regulatory network inference and clustering (SCENIC)^22,23^. Regulon activities of each cluster were broadly in line with these transcriptional changes (Figure 2I). Compared with SC0_Ctrl, SC0_PSP tended to show lower activity of IFN/IRF regulons, consistent with the downregulation of interferon-stimulated genes and cytoskeleton/adhesion pathways. In contrast, regulons linked to lipid and cholesterol metabolism (SREBF2), nuclear-encoded mitochondrial regulation (YY1 and GABP) and stress-responsive factors (ATF2 and BCLAF1) were relatively higher activity in SC0_PSP than in SC0_Ctrl , in agreement with the GSEA enrichment for protein-folding and ER-stress pathways. Notably, homeostatic G-protein–coupled receptors (*CX3CR1*, *P2RY13*, and *GPR34*) remained comparatively highly expressed despite reduced ETS activity, raising the possibility that core homeostatic identity is maintained by ETS-independent mechanisms. Taken together, these observations suggest that SC0_PSP represents a homeostatic-like microglial state that has undergone adaptive proteostatic and metabolic remodeling with enhanced MHC class II–mediated antigen presentation, rather than a strongly interferon-driven inflammatory phenotype.

Outside SC0, differences in regulon activities between PSP and controls were generally modest. SC2 (BAMs) showed the second-largest PSP–control centroid distance in the UMAP embedding (Figure 2A and 2B; centroid distance = 0.32) and yielded 31 upregulated and 12 downregulated DEGs in pseudobulk analysis (Figure 3B and Data S7). Upregulated genes included heat-shock proteins and associated chaperones (*HSPB1*, *DNAJB1*, *HSP90AB1* and the Hsp90 co-chaperone *PTGES3*), as well as genes linked to the ubiquitin–autophagy–lysosomal pathway (the selective autophagy receptor *SQSTM1*, the deubiquitinase *USP36*, the E3 ligase *WWP1*, the autophagy-regulating transcription factor *FOXO3* and the lysosomal glycoprotein prosaposin, *PSAP*). Consistent with SC0_PSP, pathway analysis detected several protein-folding–related terms (for example GO:BP chaperone-mediated protein folding), whereas clear enrichment of ER-stress/UPR pathways was not observed (Figure 3F and Data S7). In contrast, SC1, SC3 and SC4 showed only limited PSP–control differences, with 2, 11 and 0 DEGs, respectively (Figure 3A, 3C, 3D, and Data S7), and their pathway analyses were dominated by negatively enriched terms (Figure 3E, 3G, 3H, and Data S7). Regarding regulon activity, PSP samples across all subclusters (SC1–SC4) showed a reduction in SPI1/ETS homeostatic tone (Figure 3I–L). Stress-related regulons were most pronounced in SC2, clearly elevated in SC3, biased toward a proliferative/repair program in SC4, and minimal in SC1 (Figure 3I–L).

**Figure 3.**
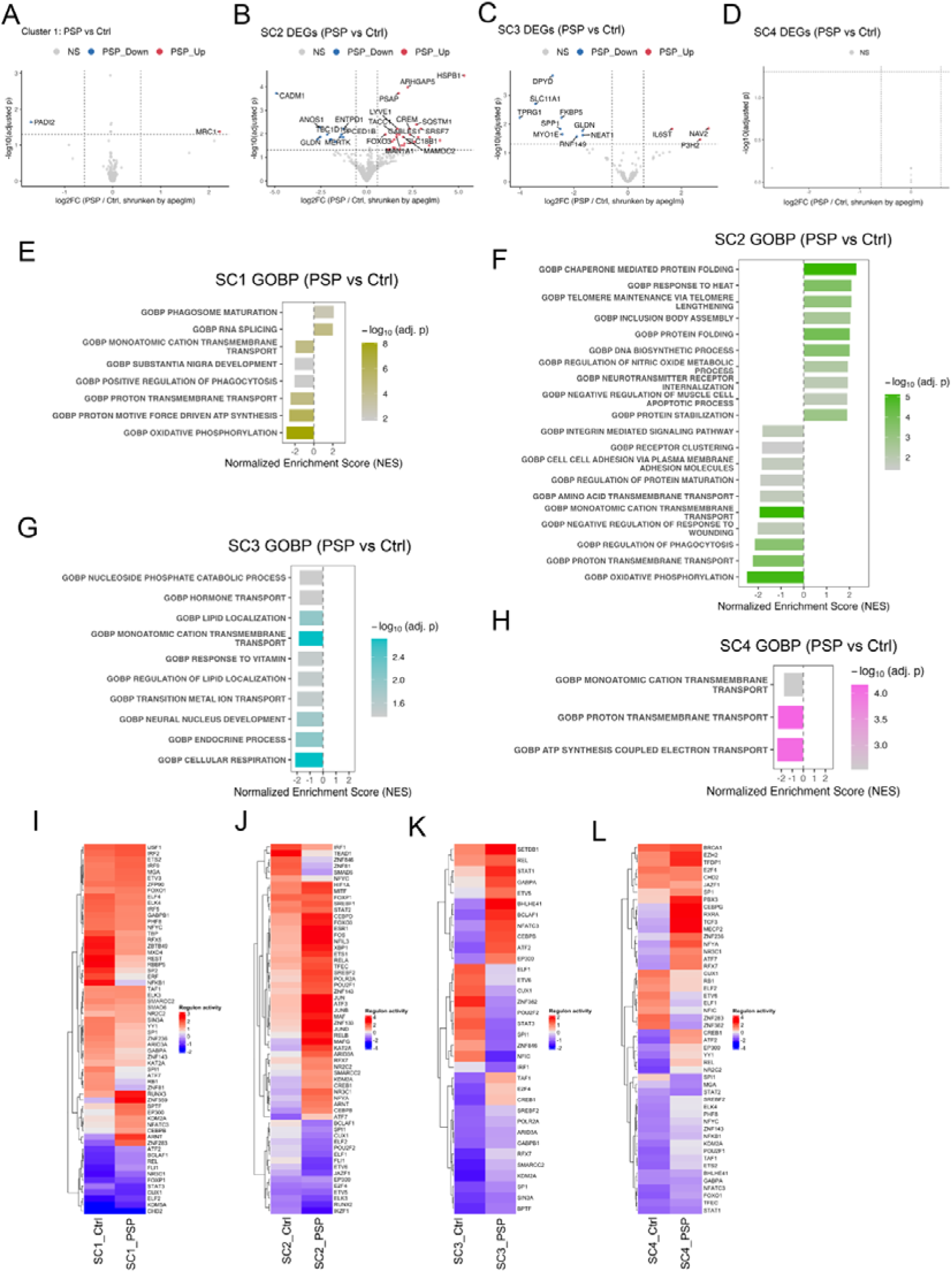
**Distinct transcriptional and regulon states of Microglia–PVM subclusters in PSP** (A-D) Differential expression between PSP and controls within Microglia–PVM subclusters SC1–SC4. Volcano plots show shrinkage-estimated loglFC (PSP/Ctrl, apeglm) versus –logll(adjusted p) from pseudobulk DESeq2 analysis. Red and blue dots indicate genes significantly up- and downregulated in PSP (adj. p < 0.05 and |loglFC| > 0.25; Benjamini–Hochberg correction). (E-H) Gene Ontology Biological Process (GOBP) enrichment for SC1–SC4 (PSP vs Ctrl). Gene-set enrichment analysis was performed on DESeq2 pseudobulk statistics using the MSigDB C5 GOBP collection. For each subcluster, bar plots show the 10 most significantly enriched pathways (adj. p < 0.05), ordered by normalized enrichment score (NES); color encodes –logll(adjusted p). (I-L) SCENIC-inferred transcription factor regulon activity in SC1–SC4 (PSP vs Ctrl). Heatmaps show regulon activity per subcluster–condition combination. Activity was quantified by AUCell AUC and z-scored within each regulon; red and blue indicate relatively higher and lower activity, respectively. Dendrograms depict hierarchical clustering of regulons and of PSP/Ctrl profiles within each subcluster. Panels: SC1 (A, E, and I), SC2 (B, F, and J), SC3 (C, G, and K), SC4 (D, H, and L).

### Microglia in the PSP Frontal Cortex Show a Shift Toward Ramified Morphology

Microglial morphology often tracks transcriptional state: in DAM-like responses, a transition from a ramified (process-bearing) to an amoeboid morphology is typically accompanied by loss of homeostatic markers. In our transcriptomic analysis, however, the most abundant PSP cluster, SC0_PSP, showed higher expression—relative to controls—of canonical homeostatic microglial genes that are typically associated with ramified, process-bearing states. We therefore investigated whether PSP microglia in vivo exhibit corresponding morphological features by performing quantitative morphometric analyses on neuropathology specimens.

IBA1 immunohistochemistry, which clearly delineates microglial processes, was performed on FFPE frontal cortex sections from eight PSP patients and eight controls (Figure 4A). Using the HALO® Microglia Activation Module, we quantified process length, process area, cell body area and the number of branch points per cell in both cortex and white matter (Figure 4B). All raw data are provided in Data S8. In total, 108,885 microglia were identified in the cortex and 73,119 in the white matter of PSP cases, compared with 99,664 and 61,475, respectively, in controls. The density of IBA1-positive microglia per unit area did not differ significantly between groups in the cortex and was only modestly higher in PSP in the white matter (Figure 4C). The process morphology differed markedly between groups. In the cortex, PSP microglia showed a a shift toward longer process lengths, resulting in a higher mean length than controls (47.8 ± 7.6 μm vs 37.0 ± 7.1 μm; Figure 4D). The effect was even more pronounced in white matter (52.8 ± 7.8 μm vs 39.6 ± 9.8 μm; Figure 4E). Process area showed a similar pattern, with a significant increase in PSP in white matter (Figure 4F and 4G). Cell body size did not differ significantly between groups in either region (Figure 4H and 4I). The number of branch points per cell was higher in PSP than in controls in both cortex and white matter (Figure 4J and 4k). Overall, microglia in the PSP frontal cortex tended to exhibit a more ramified, process-bearing morphology, with on average longer and more highly branched processes.

**Figure 4.**
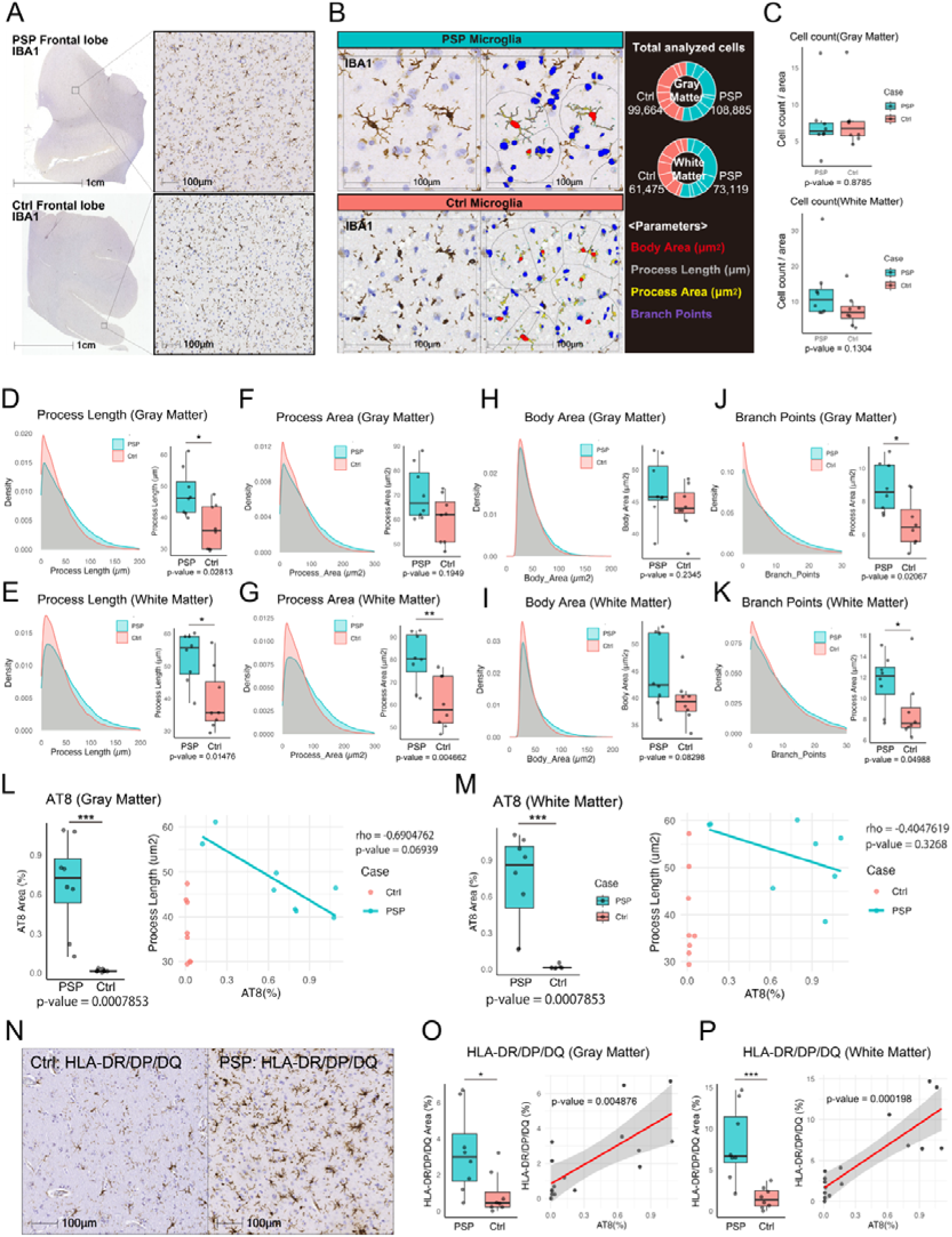
**Morphological analysis of microglia in the PSP frontal lobe** (A) Representative low- and medium-magnification images of IBA1 immunostaining in formalin-fixed, paraffin-embedded (FFPE) frontal lobe tissue from a patient with PSP and a control case. Scale bars, 1 cm (left) and 100 µm (right). (B) Example of microglial morphometric analysis using the HALO Microglia Activation Module. Quantified parameters include total process length (µm), cell body area (µm²), total process area (µm²), and number of branch points. The total numbers of analyzed cells in gray and white matter are shown for PSP and control groups. Scale bars, 100 µm. (C) Box plots of IBA1-positive microglial density in gray matter (top) and white matter (bottom). Cell counts were normalized by analyzed area (cells per area) for each case; p-values indicate group comparisons between PSP (n = 8) and controls (n = 8), with no significant differences detected. (D-K) Morphometric analysis of IBA1-positive microglia in gray matter (top panels) and white matter (bottom panels). Analyzed parameters include process length (D and E), process area (F and G), cell body area (H and I), and number of branch points (J and K). For each parameter, density plots (left) summarize all segmented cells, and box plots (right) show per-case mean values for PSP and control cases (n = 8 per group). p-values were calculated using the Wilcoxon rank-sum test (*p < 0.05, **p < 0.01). (L-M) Relationship between tau pathology and microglial process length in gray matter (L) and white matter (M). Left panels show box plots of AT8-positive area (%) per case in PSP and control groups; p-values from Wilcoxon rank-sum tests with normal approximation because of ties are indicated (***p < 0.001 for both regions). Right panels show scatter plots relating AT8-positive area (%) to mean microglial process length (µm) per case. Lines represent Spearman’s rank correlations calculated in PSP cases only (gray matter: ρ = –0.69, p = 0.069; white matter: ρ = –0.40, p = 0.33). (N-P) Quantification of HLA-DR/DP/DQ–positive microglial area and its relationship to tau pathology in PSP and control cases. (N) Representative HLA-DR/DP/DQ immunostaining in the frontal cortex. (O and P) Left, box plots showing the percentage of HLA-DR/DP/DQ–positive area in gray matter (O) and white matter (P) per case in PSP (n = 8) and control (n = 8) brains (*p < 0.05 in gray matter; ***p < 0.001 in white matter; Wilcoxon rank-sum test). Right, scatter plots illustrating the relationship between AT8-positive area (%) and HLA-DR/DP/DQ–positive area (%) per case. Correlations were assessed using Spearman’s rank correlation test, and red lines indicate linear regression fits with 95% confidence intervals (gray matter: p = 0.0049; white matter: p = 0.00020).

The AT8 positive area, a proxy for phospho-tau burden, tended to correlate negatively with mean process length at the case level, although these correlations did not reach statistical significance (Figure 4L and 4M). Given the increased expression of antigen-processing genes in SC0_PSP (Figure 2G), we also performed HLA-DR immunohistochemistry with HALO-based quantification (Figure 4N). The fraction of HLA-DR–positive microglia per unit area was significantly higher in PSP than in controls in both cortex and white matter, and this HLA-DR burden correlated positively with AT8 pathology (Figure 4O and 4P). While not establishing causality, these morphological and IHC findings are consistent with the homeostatic yet MHC class II–upregulated transcriptomic profile observed in SC0_PSP.

### Protein-folding/ER stress promotes SC0_PSP-like features in human microglia in vitro

To explore factors that might contribute to SC0_PSP-like features in the PSP frontal cortex, we performed in vitro assays using the human microglial cell line HMC3 (Figure 5A). Motivated by our pathway analyses, which showed prominent enrichment of protein-folding pathways and increased ER-stress/unfolded protein response (UPR) signatures (Figure 2D, S4A, and S4B), we used tunicamycin, an ER-stress inducer that blocks initiation of N-linked glycosylation, to increase folding load and assess phenotypic changes. Tunicamycin exhibited time- and dose-dependent toxicity in HMC3 cells: cell counts decreased after 48 h at ≥0.5 µg/mL (Figure 5B and Data S9). LDH release increased at ≥1 µg/mL after 24 h, showed a mild rise at 0.1 µg/mL with ≥48 h exposure, and rose markedly at ≥0.5 µg/mL by 48 h (Figure 5C and Data S9). Based on these data, subsequent experiments were conducted at ≤0.1 µg/mL. At 0.1 µg/mL for 48 h, ER-stress markers ATF4 and its target GADD34, as well as the ER chaperone HSP90B1, were significantly upregulated, consistent with activation of the PERK–eIF2α–ATF4 branch of the UPR suggested by the snRNA-seq data (Figure 5D and Data S9). The cytosolic chaperone HSPA1A also increased, indicating heightened folding stress originating from the ER. Collectively, these findings indicate that our HMC3 model establishes a tunicamycin-induced protein-folding/ER-stress condition with induction of a chaperone program in human microglia.

**Figure 5.**
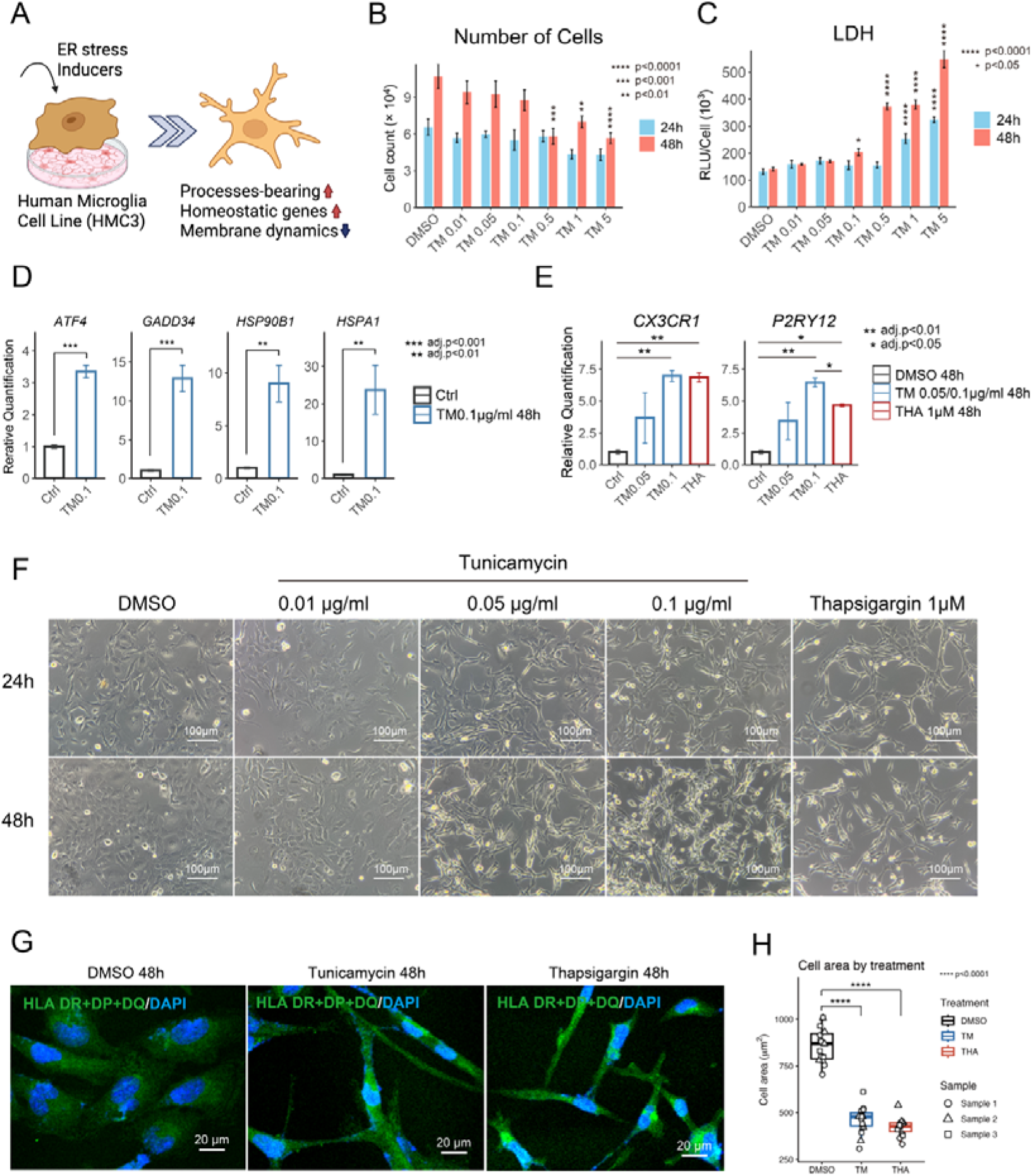
**Changes in Human microglia cell (HMC3) Associated with ER Stress** (A) Schematic graphs of in vitro assay. Created in BioRender. (B) Effects of tunicamycin on HMC3 cell numbers. Cell numbers were quantified with a Countess automated cell counter 24 h and 48 h after treatment with DMSO or the indicated concentrations of tunicamycin (n = 3 per condition). Bars represent mean ± SEM. Asterisks denote significant differences from the DMSO control at the corresponding time point, based on a linear model including Time and Treatment with Dunnett-adjusted contrasts versus the DMSO control (**p < 0.01, ***p < 0.001, ****p < 0.0001). (C) Effects of tunicamycin on LDH release. LDH activity in culture supernatants was measured 24 h and 48 h after treatment, and relative luminescence units (RLU) were normalized to cell number in each well (n = 3). Bars indicate mean ± SEM. Asterisks indicate significant differences from the DMSO control at the same time point, assessed using the same Dunnett-adjusted linear model (*p < 0.05, ****p < 0.0001). (D) Quantitative PCR analysis of protein folding / ER stress–related genes. Relative mRNA levels of *ATF4*, *GADD34*, *HSP90B1* and *HSPA1* in HMC3 cells treated with DMSO 48 h or tunicamycin 0.1 µg/mL for 48 h (n = 3; mean ± SEM). Two-sided Welch’s t-test were performed for each gene, and p values were adjusted for multiple testing across genes using the Benjamini–Hochberg correction (**adj. p < 0.01, ***adj. p < 0.001). (E) qPCR analysis of homeostatic microglial genes *CX3CR1* and *P2RY12* in HMC3 cells after 48 h treatment with DMSO, tunicamycin (0.05 or 0.1 µg/mL), or thapsigargin (1 µM) (n = 3 per condition). Data are shown as relative quantification normalized to the DMSO control (mean ± SEM). Asterisks indicate pairwise differences from the DMSO control based on two-sided Welch’s t-test with Benjamini–Hochberg correction for multiple comparisons (*adj. p < 0.05, **adj. p < 0.01). (F) Bright-field images of HMC3 cells treated with DMSO or tunicamycin (0.01, 0.05, 0.1 µg/mL) and thapsigargin (1 µM) for 24 h or 48 h. Scale bars, 100 µm. (G) Representative fluorescence images of HMC3 cells stained for HLA-DR/DP/DQ (green) and DAPI (blue) after 48 h treatment with DMSO, tunicamycin (0.1 µg/mL), or thapsigargin (1 µM). Scale bars, 20 µm. (H) Quantification of HLA-DR/DP/DQ–positive cell body area in HMC3 cells. Cell body area (µm²) was measured from thresholded HLA-DR/DP/DQ–positive regions in Fiji. Each point represents one field of view (5 fields per dish, 3 independent dishes per condition); boxes indicate the interquartile range with median, and whiskers show the range. Group differences among DMSO, tunicamycin (TM) and thapsigargin (THA) were assessed by one-way ANOVA followed by Tukey’s multiple comparison test (****p < 0.0001).

We next examined homeostatic, process-associated genes (*CX3CR1* and *P2RY12*) by qPCR. Tunicamycin treatment significantly increased both transcripts, with approximately 6–7-fold induction at 0.1 µg/mL (Figure 5E and Data S9). To probe an additional ER-stress modality, we used thapsigargin, which inhibits SERCA and depletes ER Ca²l to induce protein-folding stress; similar upregulation of *CX3CR1* and *P2RY12* was observed (Figure 5E and Data S9). These changes parallel the elevated expression of homeostatic GPCRs seen in SC0_PSP. Under DMSO control conditions, HMC3 microglia exhibited an amoeboid-like morphology with widely spread cytoplasm (Figure 5F). In contrast, exposure to protein-folding/ER stress (tunicamycin or thapsigargin) induced a dose-dependent shift toward a more process-bearing, ramified morphology: tunicamycin at 0.01 µg/mL produced no clear changes at 24–48 h, whereas concentrations ≥0.05 µg/mL (and corresponding thapsigargin treatments) yielded cells with narrow bodies and elongated protrusions (Figure 5F and 5G). Quantification of cytoplasmic area per cell (HLA-DR/DP/DQ–positive area) confirmed this morphological shift, with ER-stress–treated cells showing a significant decrease in cytoplasmic area compared with DMSO controls (Figure 5H and Data S9). In time-lapse imaging, DMSO-treated HMC3 cells typically showed an amoeboid-like morphology with prominent, rapidly fluctuating membrane ruffles. Following tunicamycin exposure, a larger fraction of cells adopted narrow, process-bearing morphologies with more stable processes and visibly reduced membrane ruffling (Video S1). Although these changes were not quantified, they are qualitatively consistent with a reduction in dynamic membrane remodeling. This pattern parallels the SC0_PSP pathway analysis, in which Rho GTPase cycle and motility-related pathways were downregulated (Figure 2D) and suggests that protein-folding / ER stress can reproduce aspects of the altered motility state inferred from the snRNA-seq data, even if it does not fully recapitulate the SC0_PSP phenotype.

### Attenuated cell–cell communication signatures in PSP microglia

In neurodegeneration, not only aberrant proteins but also cell–cell communication can influence microglial transcriptional states. To probe upstream cues that might contribute to the SC0_PSP state, we analyzed intercellular communication with CellChat^24,25^ and NicheNet^26,27^. CellChat infers intercellular communication networks from single-cell expression using a curated ligand–receptor knowledge base, whereas NicheNet prioritizes upstream ligands whose predicted target programs best explain receiver DEGs.

Using CellChat, we first compared the total number of inferred interactions and the aggregate interaction strength across all cell types. The number of inferred interactions increased from 2,756 in controls to 3,541 in PSP, whereas the aggregate interaction strength decreased from 131.204 (control) to 117.653 (PSP) (Figure 6A). These results suggest a rewiring of the communication network in PSP—more inferred edges but lower overall signaling strength. In the circle plot, all microglial subclusters except SC1 showed decreases in both the number of inferred interactions and the aggregate interaction strength (Figure 6B). In SC0_PSP, CellChat predicted fewer ligand–receptor interactions and lower overall communication strength; concordantly, GSEA highlighted reductions in Reactome pathways related to cell–cell communication and signaling (Figure S4A). Several edge genes—including *NCK2*, *DST*, *CADM1* and *ACTN1*—encode membrane scaffolds that organize receptor clustering and actin-linked adhesion; their reduced expression may contribute to the observed contraction of the cell–cell signaling network.

**Figure 6.**
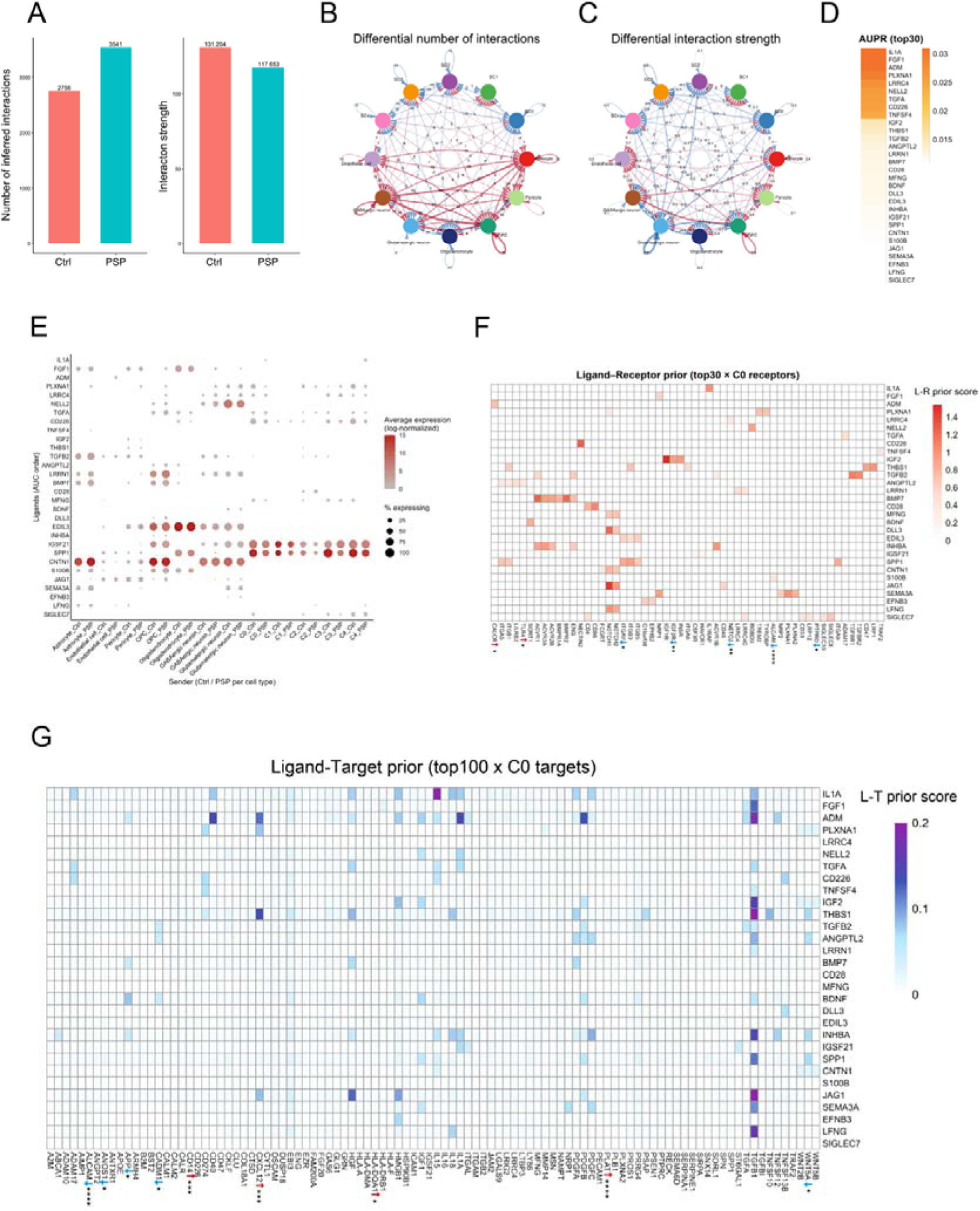
**Cell― Cell Communication** (A) Global cell–cell communication inferred by CellChat. Left, total number of significant ligand–receptor interactions per condition. Right, overall interaction strength (sum of CellChat edge weights). (B and C) Circle plots of differential cell–cell communication between control and PSP. Nodes represent cell types and edges represent ligand–receptor interactions, including self-loops for autocrine signaling. Edge color indicates the condition with stronger signaling (red, PSP > Ctrl; blue, Ctrl > PSP), and edge width reflects the magnitude of the difference. (B) Difference in the number of interactions; (C) difference in overall interaction strength. (D) NicheNet ligand activity ranking for the SC0_PSP upregulated gene set. Heatmap showing the top 30 candidate ligands ranked by ligand activity score (AUPR, area under the precision–recall curve); color indicates AUPR. (E) Expression of candidate ligands across sender cell types. Columns show sender cell types in Ctrl and PSP; rows list the top-ranked ligands. Color indicates mean log-normalized expression and dot size the percentage of expressing cells. Pseudobulk DESeq2 analysis detected no significant PSP–control differences for these ligands. (F) NicheNet ligand–receptor prior with receptor differential expression in SC0. Heatmap showing NicheNet ligand–receptor prior scores (color scale) for the top 30 ligands (rows) and receptors detected in SC0 microglia (columns). Arrows and asterisks mark receptors that are differentially expressed between PSP and control within SC0 based on pseudobulk DESeq2 analysis (Wald test with Benjamini–Hochberg–adjusted p values across all genes; *adj. p < 0.05, **adj. p < 0.01, ****adj. p < 0.0001). (G) NicheNet ligand–target prior in SC0 microglia. Heatmap of NicheNet ligand–target prior scores for the top 30 ligands (rows; ranked by AUPR) and the top 100 predicted target genes in SC0 (columns). Color indicates the ligand–target prior (regulatory-potential) score. Column labels denote differential expression of each target in SC0 (PSP vs control) based on pseudobulk DESeq2 analysis (Wald test with Benjamini–Hochberg–adjusted *p* values across all genes; red, higher in PSP; blue, lower in PSP; *adj. p < 0.05, ***adj. p < 0.001, ****adj. p < 0.0001).

We next used NicheNet to prioritize upstream cues that could induce SC0_PSP-like transcriptional changes. The target set comprised 76 upregulated DEGs (pseudobulk DESeq2). Ligand activity was quantified by the area under the precision–recall curve (AUPR) for recovery of the SC0_PSP target set, and the top-30 AUPR-ranked ligands grouped into functional categories: inflammatory mediators (*IL1A*, *ANGPTL2*, *SPP1*, *S100B*); anti-inflammatory/repair and neurotrophic cues (TGF-β/Activin/BMP, *ADM*, *IGF2*, *BDNF*, *FGF1*); integrin-associated ligands and matrix proteins for phagocytosis/motility (*THBS1*, *EDIL3*, *SPP1*); contact-dependent guidance signals (*SEMA3A*, *EFNB3*, *CNTN1*); Notch pathway components (ligand *JAG1*; *DLL3* as a cis-inhibitory component; *MFNG*/*LFNG* as Fringe modifiers); and T/NK co-stimulatory and checkpoint-like axes (for example *TNFSF4*/OX40L, with pathway partners such as *CD28*, *CD226* and *SIGLEC7*). When ligand expression was compared by sender cell type between PSP and controls (pseudobulk), no ligand showed a significant difference (Figure 6C and Data S10). By contrast, receiver-side receptors in SC0_PSP displayed selective shifts (Figure 6D): up (*CALCR*, *TLR4*) and down (*ALCAM*, *IGF2R*, *NETO2*, *ITGAV*, *PTPRO*). The heat map shows LR_prior values between the top-30 AUPR ligands and receptors detected in ≥5% of SC0 (Homeostatic) cells (Figure 6D and Data S10). Together, these data point to receiver-side tuning in SC0_PSP rather than broad changes in ligand supply.

We next examined downstream target expression associated with the prioritized ligand–receptor (L–R) signals. For the *SPP1*–integrin axis, reduced *ITGAV* in SC0_PSP could attenuate *SPP1*–*ITGAV* signaling; consistent with this, *WNT5A*—a NicheNet-predicted target of this axis—was decreased, and *SPP1* (primarily released by SC0 itself) showed a non-significant trend toward lower expression in SC0_PSP (Figure 6E and Data S10). Given that *SPP1* is a DAM-associated mediator implicated in inflammation and motility, reduced *SPP1*–integrin signaling may be compatible with maintenance of a more homeostatic program in SC0_PSP. For *TLR4*, the receptor was upregulated in PSP, whereas *WNT5A* and *CADM1*—predicted downstream genes within an *ANGPTL2*-implicated pathway—were decreased. *ANGPTL2* was mainly expressed by oligodendrocytes and tended to be lower in PSP (Data S10), which could be consistent with receiver-side compensation (receptor upregulation despite reduced ligand supply).

Finally, *CALCR* was upregulated in SC0_PSP. Although *ADM* counts were low, it tended to be higher in astrocytes, endothelial cells, GABAergic neurons and pericytes in PSP. NicheNet implicated an *ADM*–receptor axis involving *CALCR*, and the predicted target *CXCL12* was increased; by contrast, *APP* decreased, suggesting contributions from additional cues (for example *BDNF* and *BMP7*) within this network context.

Taken together, overall inferred communication strength was reduced, and only a limited subset of ligand–receptor edges showed clear correspondence with the upregulated DEG set in SC0_PSP. Notably, the hallmark transcriptomic increases that define SC0_PSP were not well explained by intercellular signaling in this analysis, pointing instead toward a prominent contribution from cell-intrinsic or microenvironmental stress pathways.

## DISCUSSION

In this study, we systematically characterized microglial states in the frontal cortex of PSP and control brains using snRNA-seq, regulon analysis, in vitro assays and quantitative histopathology. Within the homeostatic microglial compartment, we identified a distinct PSP-enriched state marked by protein-folding/ER-stress signatures. This PSP-related microglial state is characterized by (i) enhanced protein-folding/ER-stress pathways, (ii) an augmented homeostatic program with a shift toward a process-bearing, ramified morphology, (iii) increased antigen presentation via MHC class II, (iv) reduced motility and cytoskeleton/adhesion pathways, (v) attenuation of IFN/IRF, and (vi) contraction of inferred cell–cell communication (Figure 7). Our in vitro experiments suggest that protein-folding/ER stress is sufficient to induce a subset of these features—such as upregulation of homeostatic genes and a transition toward a more ramified morphology with reduced membrane ruffling. To provide a concise descriptor for this constellation of changes, we refer to this PSP-related microglial state as protein-folding/ER–stress–associated microglia (PSAM). We use the term ‘PSAM’ to denote a PSP-associated microglial state characterized by protein-folding and ER stress pathways, while acknowledging that PSP microglia likely encompass additional distinct states beyond PSAM.

**Figure 7.**
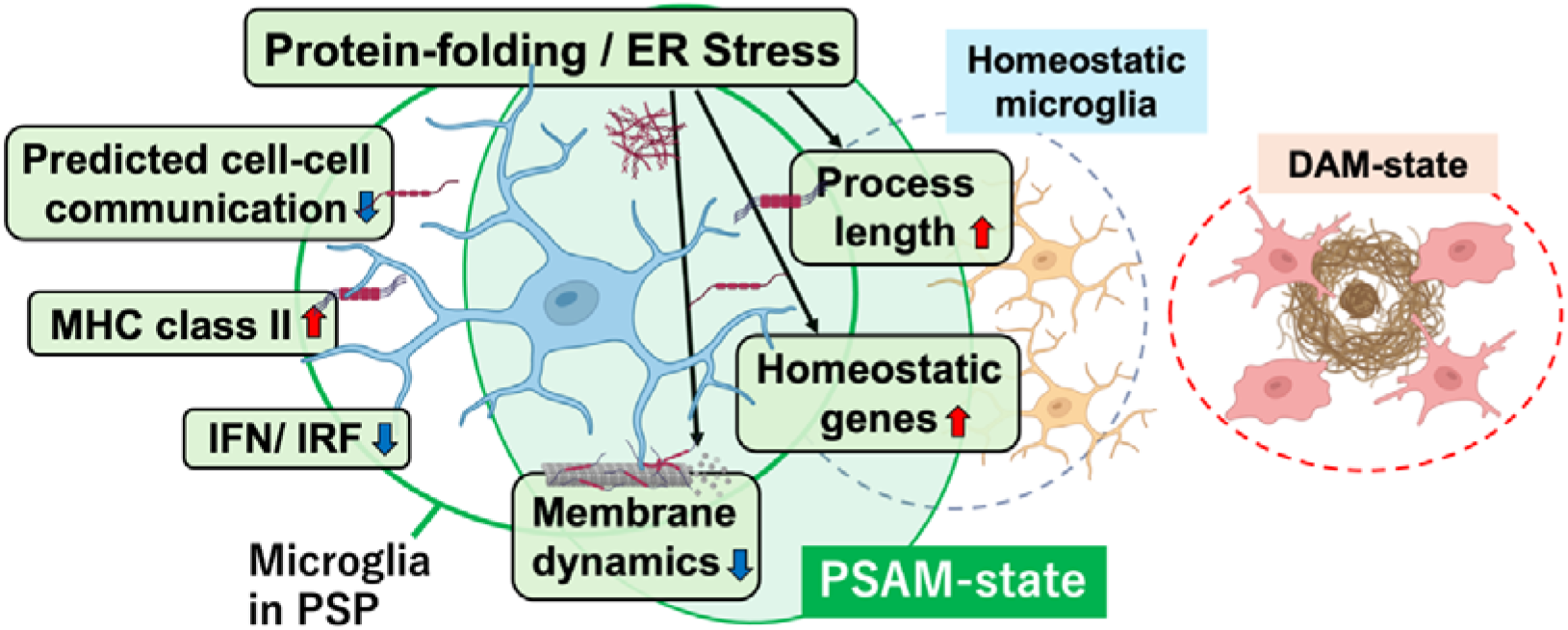
**Graphical abstract for distinct microglia in PSP frontal cortex.** Distinct microglial state identified within homeostatic microglial clusters in PSP. A protein-folding/ER-stress–associated microglial (PSAM) state, characterized by homeostatic genes and elongation of microglial processes, which exhibits properties that are largely opposite to those of classical disease-associated microglia (DAM). Created in BioRender.

Neurodegeneration-associated microglia have largely been conceptualized in terms of the DAM state, a response to amyloid plaques and dying cells characterized by a macrophage-like program enriched for phagocytosis, lipid metabolism and complement ^3–5^, with downregulation of homeostatic markers and a more amoeboid morphology. The classical activation lineage described by Del Río-Hortega^28^ aligns with this view. By contrast, PSAM accentuates, rather than extinguishes, homeostatic features. In this sense, PSAM represents a protein-folding–stress–associated, “a homeostatic-mimicking microglial state” that is phenotypically opposite to the inflammatory, phagocytic DAM pole. RNA-velocity analysis further supports the idea that PSP microglia do not lie on a single linear trajectory toward a DAM-like endpoint. Velocity vectors from SC1 (Neuronal surveillance) and SC0 (Homeostatic) toward SC2 (BAMs) are consistent with a drift toward macrophage-like, BAM/DAM-associated modules, whereas a PSP-enriched convergence point within SC0 indicates an alternative route (Figure 1I and 1J). These patterns suggest that, within the homeostatic compartment, microglia may bifurcate into two inferred trajectories: a BAM/DAM-like arm and a PSP-related arm that includes the PSAM state. We emphasize that these trajectories are inferred from cross-sectional data and do not establish lineage relationships, but they provide a useful conceptual framework for understanding coexisting, and partially opposing, microglial states in PSP.

Protein-folding stress and subsequent ER stress are strongly implicated in PSP, with GWAS repeatedly reporting EIF2AK3 (PERK) as a risk locus^29,30^. Consistently, Whitney et al. demonstrated activation of the integrated stress response (ER stress/UPR) in microglia by single-nucleus RNA-seq of PSP brains^31^. However, PERK–eIF2α–ATF4 signaling and related proteostasis pathways are not unique to PSP; rather, they constitute a recurrent stress axis across multiple neurodegenerative diseases, including Alzheimer’s disease, tauopathies, synucleinopathies and TDP-43 proteinopathies^32^. This raises the possibility that PSAM-like transcriptional states reflect a shared microglial response to chronic neurodegenerative stress across diseases, rather than a PSP-specific phenomenon. Our data support this view at least in the context of tau pathology. In gene-set clustering analysis, the tau-related AD2 microglial signature reported by Emma et al. clustered in a homeostatic module rather than with DAM-like modules (Figure 1E). Consistently, this AD2 gene set was significantly and positively enriched in a pre-ranked GSEA using the SC0_PSP versus SC0_Ctrl gene ranking (Figure 2E), suggesting that a PSAM state may be shared across tauopathies.

The expression of PSAM-like states is likely to be shaped, and in some settings masked, by the surrounding brain microenvironment. Amyloid-β alone can profoundly alter microglial function^33^, and mixed pathology with amyloid plaques may superimpose a DAM-like program on an underlying PSAM-like background ^34^. As disease progresses, microglial responses to dying cells^35,36^ may further strengthen DAM-like signatures over time, obscuring PSAM-like states in late-stage or mixed-pathology brains. In our neuropathological analysis of PSP frontal cortex samples lacking detectable amyloid-β deposition, the fraction of AT8-positive area showed a non-significant trend toward negative correlation with microglial process length. Although exploratory, this pattern is compatible with PSAM-like, ramified morphologies being most evident in tau-predominant regions with relatively preserved tissue integrity and becoming increasingly obscured as tau burden and other disease-associated signals accumulate.

Our cell–cell communication analyses are consistent with this view of a largely stress-aligned, rather than strongly ligand-driven, PSAM-like state. CellChat revealed that, across microglial subclusters, the total number and overall strength of inferred interactions were reduced in PSP, with all microglial subclusters except SC1 showing fewer and weaker outgoing and incoming edges. In SC0_PSP specifically, CellChat predicted a contraction of both interaction number and aggregate communication strength, and GSEA likewise indicated reduced cell–cell communication pathways. NicheNet analysis identified a set of candidate ligands, but pseudobulk comparisons did not reveal robust ligand-level changes, and only a modest fraction of SC0_PSP-upregulated genes could be accounted for by the top ligand–receptor pairs. Together, these findings suggest that the PSP-related transcriptional shift in SC0 is not primarily driven by large changes in ligand supply from other cell types but is more compatible with a state shaped by cell-intrinsic and local microenvironmental stressors, such as tau and proteostasis burden.

Within this context, CX3CR1 provides a potential mechanistic link between tau, preserved homeostatic features and altered signaling in PSAM. Although the canonical ligand of CX3CR1 is CX3CL1, CX3CR1 has also been implicated in the uptake of extracellular tau^37^, and increased CX3CR1 expression in PSAM could therefore facilitate tau handling. Notably, in our data, the GO:BP G protein–coupled receptor signaling pathway was downregulated despite sustained high expression of CX3CR1. In light of prior work showing that tau and CX3CL1 can compete for CX3CR1 binding^38^, one speculative interpretation is that CX3CR1 becomes increasingly engaged in atypical ligand interactions such as tau binding and uptake, rather than transmitting its canonical CX3CL1-dependent GPCR signals.

Our study has limitations, including the use of cross-sectional human tissue^39^, the focus on the frontal cortex and a modest sample size. Microglia exhibit region-specific heterogeneity ^39^, and it will be important to determine whether PSAM-like states generalize across other PSP-affected regions and across additional neurodegenerative cohorts. Moreover, our in vitro experiments used an immortalized human microglial cell line and pharmacological ER-stress inducers, which capture selected aspects of PSAM but do not fully recapitulate the *in vivo* milieu. Despite these limitations, a key strength of our work is its multidisciplinary approach integrating post-mortem neuropathology with transcriptomic findings and mechanistic validation in a cell culture model.

Our findings challenge the DAM-centric paradigm and introduce a tauopathy-linked, plaque-independent microglial program with implications for therapeutic strategies targeting early tau-driven disease stages. Defining PSAM underscores that microglial responses in neurodegeneration are not limited to loss of homeostatic identity and adoption of a DAM program, but can also involve a stressed, homeostatic-like state driven by proteostasis pathways.

## RESOURCE AVAILABILITY

### Lead contact

Requests for further information and resources should be directed to and will be fulfilled by the lead contact, Gabor G Kovacs or Satoshi Tanikawa.

### Materials availability

This study did not generate new unique reagents.

### Data and code availability

Raw single-nucleus RNA-seq sequencing data generated in this study have been deposited in the NCBI Sequence Read Archive (SRA), and processed/annotated data have been deposited in the NCBI Gene Expression Omnibus (GEO). Accession numbers are pending and will be released publicly upon journal publication. Custom code used for preprocessing and downstream analyses is available from the corresponding author upon reasonable request.

## ACKNOWLEDGMENTS

We thank Ali Mohammadi Karakani and Jun Li for assistance with specimen preparation, antibody optimization, and immunostaining. We thank Helen Chasiotis for assistance with nuclei extraction from brain tissue and library preparation. We also thank the Princess Margaret Genomics Centre for sequencing services. This work was supported by the Edmond J. Safra Philanthropic Foundation, Parks Foundation and the Rossy Family Foundation. Additional support was provided by the Laura Sabia Research Awardee.

## AUTHOR CONTRIBUTIONS

S.Tanikawa conceived the study, analyzed the snRNA-seq data, performed microglial morphometric analyses, carried out *in vitro* experiments, prepared the figures and wrote the manuscript. T.K. contributed to snRNA-seq data analysis. S.Tanaka. critically reviewed and revised the manuscript. G.G.K. provided overall supervision, experimental guidance, obtained funding and revised the manuscript. All authors read and approved the final manuscript.

## DECLARATION OF INTERESTS

The authors declare no competing interests.

## DECLARATION OF GENERATIVE AI AND AI-ASSISTED TECHNOLOGIES

During the preparation of this work, ChatGPT (OpenAI) was used to assist in drafting and refining analysis code (e.g., code structure and troubleshooting). After using this tool, the author reviewed and edited the content as needed and takes full responsibility for the content of the publication.

## SUPPLEMENTAL INFORMATION

**Document S1: Figures S1–S4**

**Data S1-10: Source data for Figure S1–S4 and Figure 1-6.**

**Video S1: Effect of tunicamycin on HMC3 microglial motility**

## STAR METHODS

### KEY RESOURCES TABLE

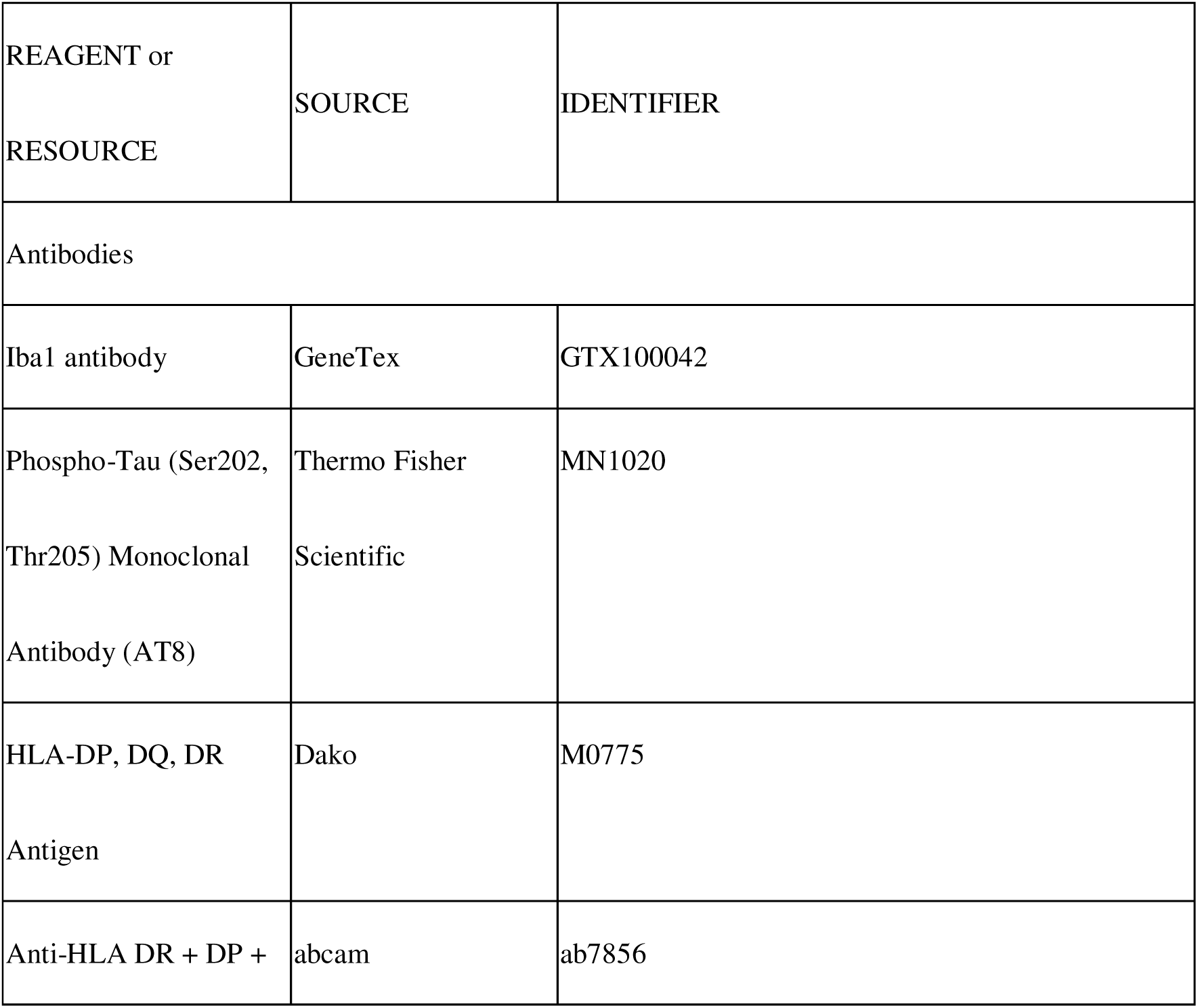
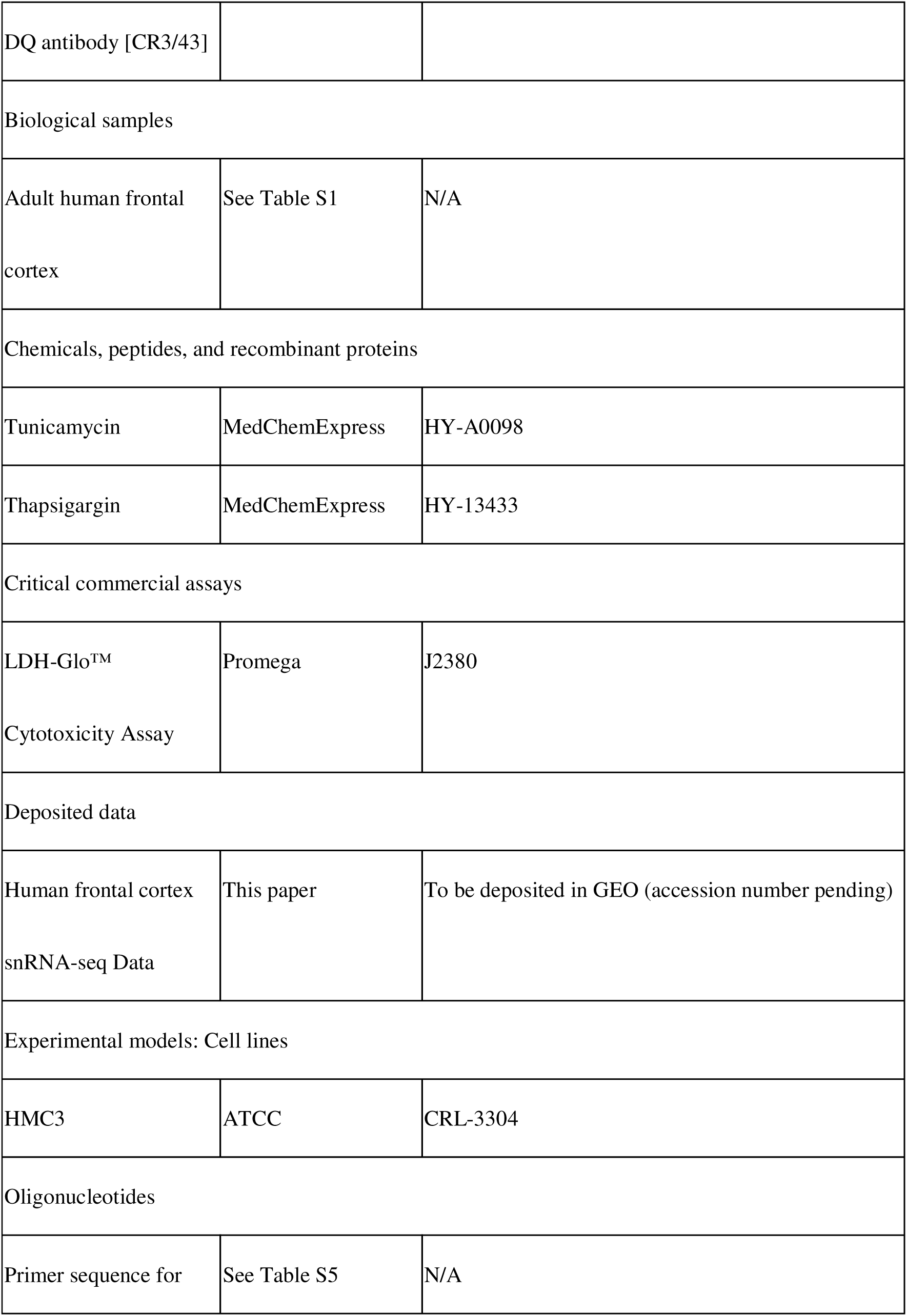
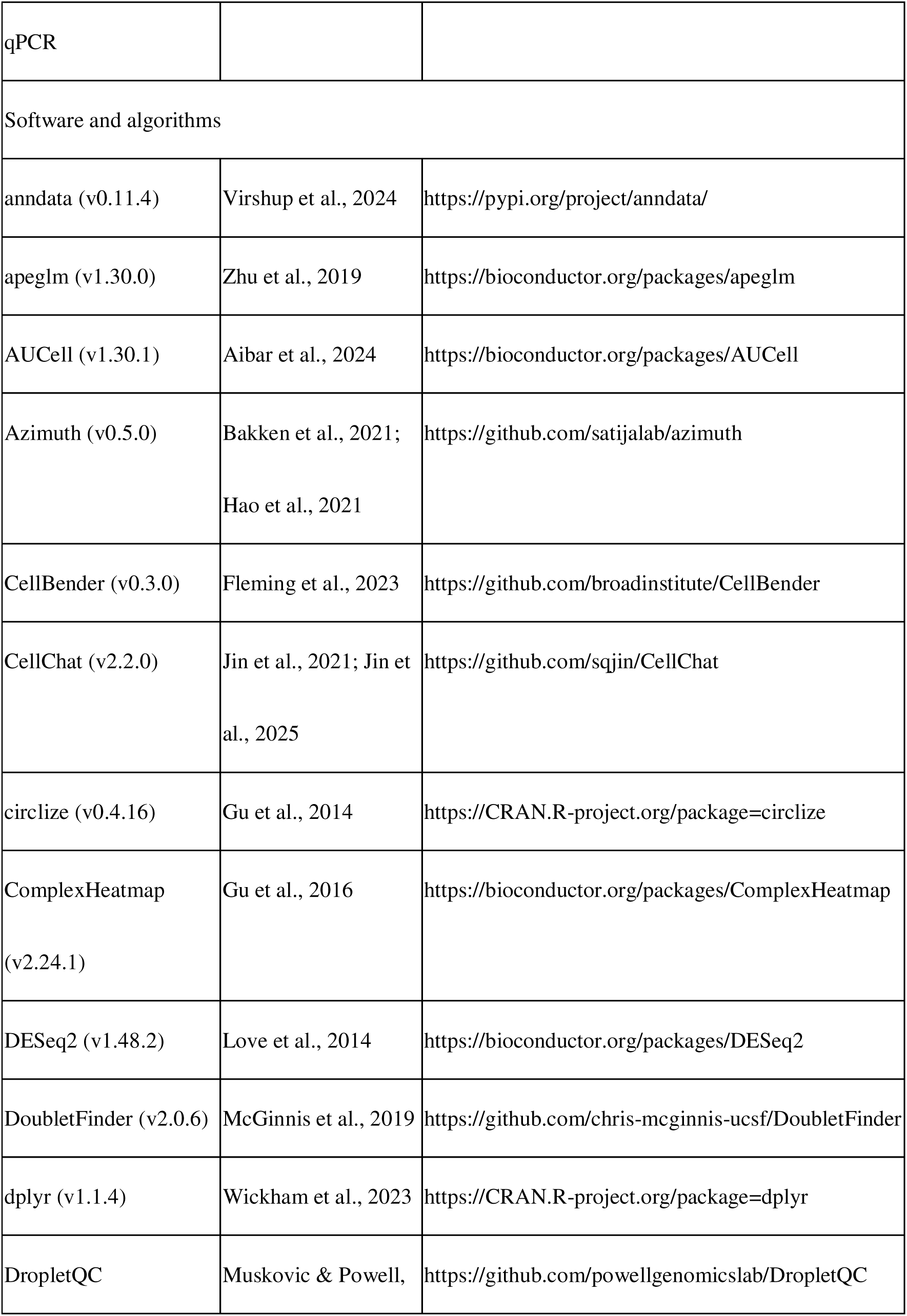
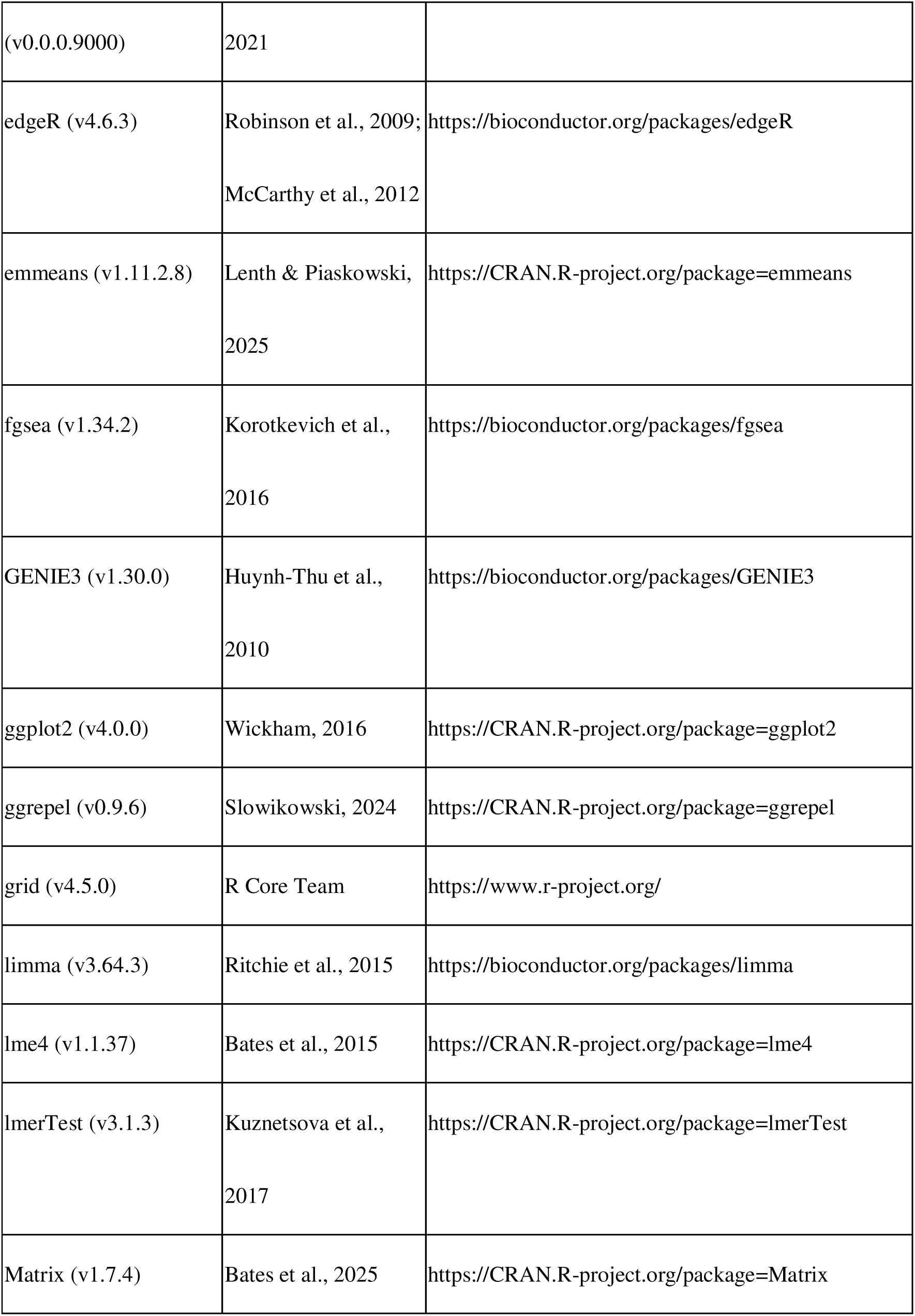
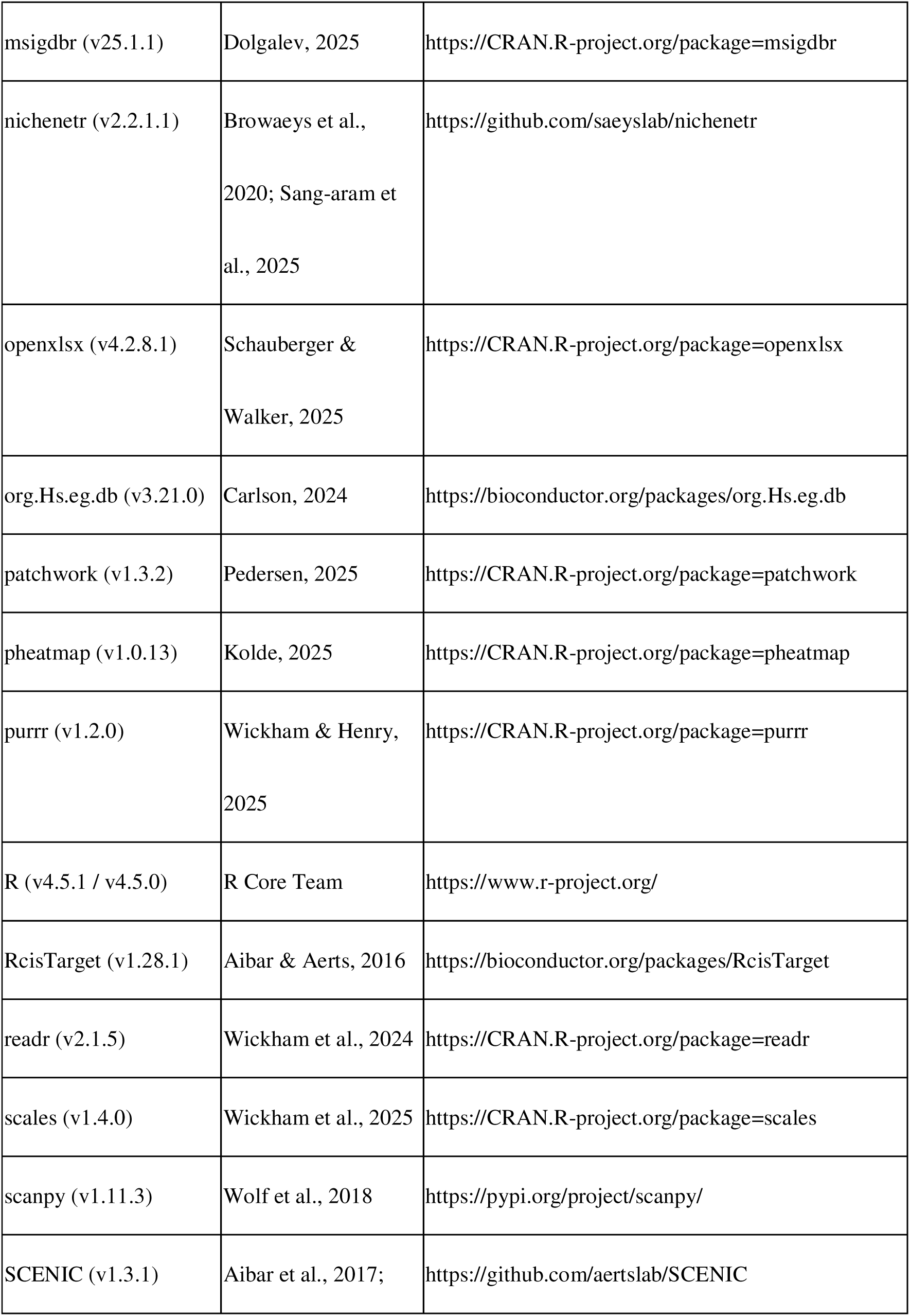
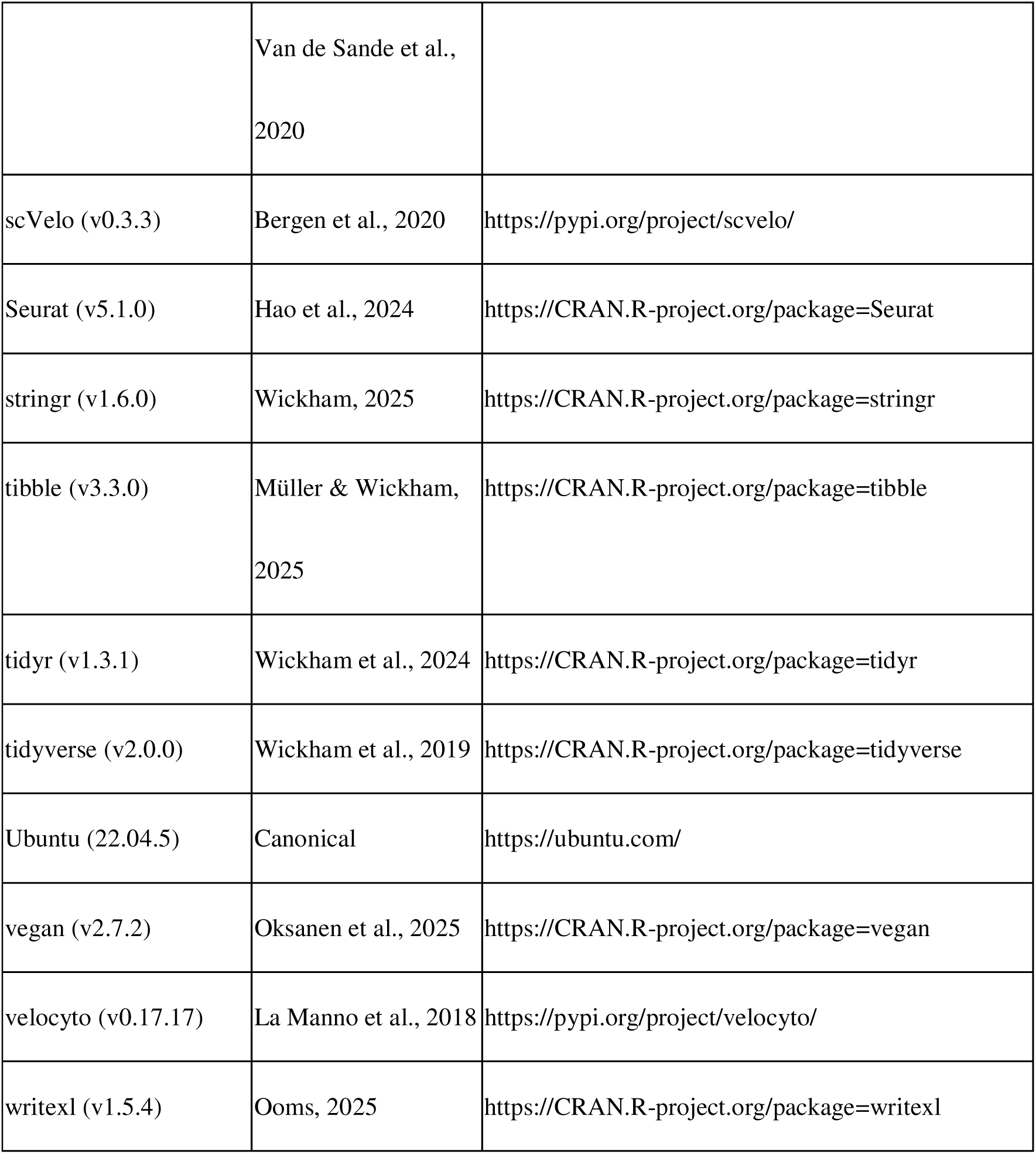

## METHOD DETAILS

### Human brain samples, case selection, and processing

Human autopsy brain tissue was collected with informed consent and ethics approval from the University Health Network Research Ethics Board (Nr. 20-5258). Cases were selected from the University Health Network–Neurodegenerative Brain Collection (UHN–NBC) based on definitive neuropathological diagnosis and systematic assessment of neuropathologies ^40,41^.

Cases diagnosed with PSP based on neuropathology and lacking co-pathology were selected. Age-matched control cases were also selected. All samples were derived from the middle frontal. At autopsy, one hemisphere was coronally sectioned, immediately flash-frozen, and stored at −80 °C. For snRNA-seq, we analyzed 8 PSP and 5 control cases using fresh-frozen tissue; for immunohistochemistry (IHC), we examined 8 PSP and 8 control cases using formalin-fixed paraffin-embedded (FFPE) tissue. Case information is provided in Data S1.

### Nuclei Isolation, Library Preparation, and Sequencing for snRNA sequencing

Frozen frontal cortex tissue was processed using two nuclei-isolation workflows. For PSP2–4 and Control1–3, nuclei were isolated on a Singulator 100 (S2 Genomics, Livermore, California, USA) following the manufacturer’s instructions. For PSP1 and PSP5–8 and Control4–5, nuclei were isolated using the Chromium Nuclei Isolation Kit with RNase Inhibitor (10x Genomics, Pleasanton, California, USA). For the Singulator-isolated cases (PSP2–4, Control1–3), libraries were prepared with the Chromium Next GEM Single Cell 3′ Kit v3.1 (10x Genomics, Pleasanton, California, USA). For the 10x-kit cases (PSP1 and PSP5–8, Control4–5), libraries were prepared with GEM-X Universal 3′ Gene Expression v4 (10x Genomics). The Singulator-isolated cohort (PSP2–4, Control1–3) were sequenced on an Illumina NovaSeq 6000 (Illumina, San Diego, California, USA), whereas the 10x-kit cases(PSP1 and PSP5–8, Control4–5) were sequenced on an Illumina NovaSeq X (Illumina, San Diego, California, USA). All sequencing was performed at the Princess Margaret Genomics Centre (Toronto, Ontario, Canada). Extended information is provided in Source Data S1.

### snRNA-seq data preprocessing, integration and cell annotation

Raw sequencing data were processed using the Cell Ranger pipeline (version 9.0.1; 10x Genomics). Reads were aligned to the human reference genome (GRCh38-2024-A), and because single-nucleus RNA-seq was performed, reads mapping to both exonic and intronic regions were included in the gene–barcode count matrices. Libraries prepared with the GEM-X Universal 3′ Gene Expression v4 assay were run using on-chip multiplexing (OCM): two cases were loaded into separate input channels corresponding to a single recovery well, and samples were subsequently demultiplexed based on their channel-specific Gel Bead barcodes using the cellranger multi pipeline. This yielded separate filtered feature–barcode matrices and BAM files for each case. Libraries prepared with the Chromium Next GEM Single Cell 3′ v3.1 kit were processed on a per-sample basis using the cellranger count pipeline with default parameters.

CellBender ^13^ was run with default parameters on Cell Ranger^42^ (v9.0.1) raw matrices for each case to remove empty droplets and ambient RNAs. Seurat^43^ (v5.3.0) objects were generated from CellBender-filtered counts. Cells expressing less than 200 or more than 5,000 unique features and those with a mitochondrial RNA content of 10% or more were filtered out. Data were normalized with LogNormalize function (scale.factor = 10,000; log1p), 2,000 highly variable genes were selected, and data were scaled based on these genes. Dimensionality reduction was performed by principal component analysis (PCA), followed by Uniform Manifold Approximation and Projection (UMAP)^44^. The number of principal components (PCs) was determined according to the result of elbow plot (inspected up to 50 PCs). Nearest neighbor graph construction was performed using FindNeighbors function with the first 20 PCs and then cell clustering was conducted using FindClusters function with a resolution set to 0.3. Doublets were identified with DoubletFinder package^14^. pK value was estimated using paramSweep function with the first 20 PCs. The expected doublet rate was set to 8% and homotypic adjustment was applied (other settings default). Across 13 samples (5 Ctrl, 8 PSP), per-sample preprocessing was followed by batch correction with Harmony ^45^ via IntegrateLayers(method = HarmonyIntegration). Nearest neighbor graph construction and clustering was performed with the first 30 integrated PCs (FindNeighbors; FindClusters, resolution = 1), and UMAP was computed. Cell types were assigned by reference mapping with Azimuth ^15^ (human cortex reference). For each Seurat cluster, subclass composition was computed; clusters with ≥75% of a single subclass were labeled as those types (astrocyte, oligodendrocyte, OPC, microglia/PVM, endothelial cell, pericyte, glutamatergic neuron, GABAergic neuron), and others were labeled Undetermined and excluded from cell-type–specific analyses (Extended Data Fig. 1e). A complete list of software packages and version information are provided in the STARIZMETHODS and Data S1.

### Removal of Ambient cluster

To mitigate ambient RNA, we followed Caglayan et al.,^21^ and their GitHub (https://github.com/konopkalab/Ambient_RNA_In_Brain_snRNAseq). First, Seurat objects were generated from the raw matrix of Cell Ranger outputs. The nuclear_fraction_annotation function (from DropletQC^46^ package) was used to calculate the intronic read ratio; candidate ambient clusters were detected with ambClusterFind function (scaled on 10,000 variable genes) and ambient markers were obtained with ambMarkFind function (the ambient markers are listed in Data S2). For subcluster cleaning, we performed several steps for each non-neuronal major type (astrocyte, oligodendrocyte, OPC, microglia-PVM, endothelial, pericyte). We performed subclustering (dims = 1–15; resolution = 1.0), obtained cluster marker genes (FindAllMarkers, default settings), and tested for enrichment of ambient genes using a Fisher’s exact test, and the resulting p-values were adjusted using the Benjamini–Hochberg correction. Clusters with high levels of enrichment of ambient RNAs (FDR is less than 0.001 and odds ratio is more than 3) compared to other clusters were labeled as ambient and removed from downstream analyses. After filtering, the dataset comprised 96,674 high-quality nuclei.

### Microglia-PVM subclustering

From the integrated object, the Microglia/PVM subset was extracted (after ambient-enriched clusters had been removed as described above). The RNA assay was split by sample (orig.ident) to create layers, and for each layer we ran LogNormalize function (scale factor was set to 10,000), selected highly 2,000 variable genes, applied ScaleData, function ,and performed PCA. Based on inspection of the elbow plot, the first 15 PCs was retained for downstream steps.

Batch effects across samples were corrected with Harmony via Seurat’s IntegrateLayers function (method = HarmonyIntegration), yielding integrated reduction. Nearest neighbor graph was built on the first 15 PCs (FindNeighbors) and clusters were identified with FindClusters (resolution = 0.3). UMAP was computed from the first 15 integrated PCs for visualization. This procedure yielded five microglial subclusters used in downstream analyses.

### Differential abundance analysis of cellular composition

Cell counts per donor and Seurat cluster were aggregated from the metadata (seurat_clusters, orig.ident) to build a cluster-by-sample count matrix, with total cells per sample computed for proportion analysis. Differential testing used a precision-weighted linear model implemented in the R package limma^47^ (functions lmFit, contrasts.fit and eBayes) with group as the main effect and covariates for age (z-scored), sex, and chemistry. The PSP vs Ctrl contrast was evaluated via lmFit, contrasts.fit, and eBayes. Effect sizes were reported as log odds ratios, with 95% confidence intervals obtained using topTable (confint = 0.95). p-values were controlled for multiple testing by the Benjamini–Hochberg correction with adj. p < 0.05 considered significant. Forest plots depicted odds ratios and 95% confidence intervals on a log scale, with OR = 1 as the reference.

### Differential expression (DE) analysis

Subcluster-defining markers were identified from pseudobulk RNA-seq profiles using the R package edgeR^48^. For each donor–subcluster combination, raw UMI counts were summed; donor–subcluster samples with <10 cells were discarded. Genes were filtered with filterByExpr, libraries were normalized by TMM (calcNormFactors), and a negative-binomial GLM was fitted with design ∼ 0 + subcluster + donor to account for inter-donor variability. For each subcluster, we contrasted that subcluster against the mean of all other subclusters using the quasi-likelihood F-test (glmQLFTest^49^). Genes with adj. p < 0.05 (Benjamini–Hochberg) and positive log_2_FC were defined as enriched in that subcluster. A more stringent core set was defined using TREAT (glmTreat ^50^) with lfc = log_2_(1.5), again retaining only genes with adj. p < 0.05 and positive log_2_FC. Within each Seurat subcluster, differential expression between PSP and Ctrl was tested using DESeq2^51^. For each subcluster, raw UMI counts were summed per gene per donor, retaining donors that contributed ≥10 cells to that subcluster. A sample-level metadata table included group (Ctrl/PSP), age (z-scored), sex, and chemistry. We retained genes with ≥20 counts in at least 20% of samples within either group (minimum two samples). DESeq2’s default pipeline was used to estimate size factors (median-of-ratios), dispersions, and fit negative-binomial GLMs. The Wald test was applied to the PSP vs Ctrl coefficient; log_2_FC were shrunken with apeglm^52^, and adj. p from the unshrunken Wald results were corrected by Benjamini–Hochberg correction.

### Module-score analysis

Per-cell scores were computed with Seurat using the function AddModuleScore^53^ for 32 literature-derived upregulated gene sets^16–19^ plus a neurotransmitter-receptor signature^18^. For Microglia-PVM subclusters (SC) 0–4, scores were aggregated to donor×subcluster means and analysed using a weighted linear mixed-effects model in the R package lme4^54^, of the form score ∼ seurat_clusters + (1 | orig.ident), score ∼ seurat_clusters + (1 | orig.ident), with *seurat_clusters* as a fixed effect, *orig.ident* (donor) as a random intercept and weights equal to the number of cells contributing to each donor–subcluster average. Global cluster effects were tested by ANOVA using the R package lmerTest^5355^ with Benjamini–Hochberg correction across modules. Pairwise contrasts between clusters were obtained with the R package emmeans^56^ and adjusted by the Benjamini–Hochberg method. For module-score analyses, adjusted P < 0.01 was considered significant; for the neurotransmitter-receptor signature, adjusted P < 0.05 was used.

### Centroid distance on UMAP

For microglial subclusters SC0–SC4, we summarized Ctrl–PSP separation by the Euclidean distance between group centroids in two-dimensional UMAP space using the first 15 PCs. Centroids were the mean UMAP_1/UMAP_2 coordinates of cells in each group within a subcluster; subclusters lacking either group were omitted. Distances were visualized by segments connecting the two centroids.

### Pathway enrichment analysis

Within each microglial subcluster, we performed pathway enrichment on a ranked gene list derived from the DESeq2 analysis. For a given subcluster, raw UMI counts were summed per gene per donor with group labels (Ctrl/PSP), and a DESeq2 model ∼ group, age (z-scored), sex and chemistry were fitted. Genes were retained if they had ≥4 counts in at least 20% of samples. From the PSP vs Ctrl comparison, we used the signed Wald statistic as the ranking metric and sorted genes in decreasing order. Gene sets were obtained from MSigDB^57^ collections C5: GOBP^58^ and C2: REACTOME^59^ (Homo sapiens) via the R package msigdbr^60^. Enrichment was computed using the R package fgsea^61^ (minSize = 15, maxSize = 500; default permutations). To reduce redundancy, we applied collapsePathways and report the resulting main pathways. For each pathway we report the NES, nominal P, and adj. p (Benjamini–Hochberg); unless stated otherwise, adj. p < 0.05 was considered significant. Leading-edge genes were recorded for visualization and exported along with the results.

### RNA velocity analysis

For each case, spliced and unspliced counts were quantified from Cell Ranger–aligned BAMs using the python package velocyto^62^ and loom files were generated. Expressed repeat annotation of the human genome downloaded from the UCSC genome browser was used to mask expressed repetitive elements. AnnData^63^ objects were created from the exported Seurat object data. RNA velocity and latent time were calculated using scVelo’s^20^ dynamical workflow with default parameters. Velocity fields were visualized on the UMAP.

### Transcription regulation analysis

Gene regulatory network analysis was performed using Single-Cell Regulatory Network Inference and Clustering (SCENIC^22,23^) to identify regulon activities in each microglia cluster. Starting from the Seurat object, we used the raw UMI counts as input. Co-expression networks were learned with GENIE3^64^. Motif enrichment–based pruning was performed with RcisTarget^65^ against the human cisTarget feather ranking databases (hg19, *hg19-500bp-upstream-7species.mc9nr.feather* and *hg19-tss-centered-10kb-7species.mc9nr.feather*) and the motifAnnotations_hgnc_v9 TF–motif annotation, using default RcisTarget thresholds to define high-confidence regulons. Per-cell regulon activity was quantified as AUC scores with AUCell^66^. Estimated regulons with high confidence were visualized using the ComplexHeatmap^67^.

### CellChat-based Cell–cell communication

Cell–cell interactions among SC0–SC4 and other cell types were inferred with CellChat^24,25^ (human database CellChatDB.human, including Non-protein signaling). For each condition (Ctrl, PSP) we built a CellChat object from the integrated Seurat object and ran the standard pipeline. We defined Number of inferred interactions as the count of significant LR pairs after filtering, and Interaction strength as the sum of edge weights (communication probability/information flow). Ctrl/PSP objects were merged with mergeCellChat function, and differences were summarized at network and pathway levels.

### NicheNet-based ligand prioritization and pathway enrichment

Focusing on microglia SC0 as receiver, we used NicheNet^26,27^ with its curated lr_network and ligand_target_matrix. Receiver background genes were those detected in ≥5% of SC0 cells, and the geneset of interest comprised SC0 genes upregulated in PSP vs Ctrl from donor-level pseudobulk DESeq2. Sender-expressed genes were defined per non-microglial type at ≥5% detection and unioned. Potential ligands were required to be expressed in senders, have a cognate receptor detected in SC0 (≥5%), and be present in NicheNet’s ligand universe. Ligand activity was computed with predict_ligand_activities using Area Under the Precision–Recall curve (AUPR); the top 30 ligands were visualized. For expression evidence, sender-side ligand dot plots (mean log-normalized expression; % expressing) were shown and PSP–Ctrl differences were tested per sender by pseudobulk DESeq2. Receptor evidence in SC0 was summarized as an LR-prior heatmap restricted to receptors detected in SC0 (≥5%) with DESeq2 annotations. For ligand-centered pathway analysis, each ligand’s gene set was defined as the top 200 positively weighted targets from the ligand_target_matrix. Genes were preranked by the unshrunken signed DESeq2 Wald statistic (PSP vs Ctrl in the receiver). We ran fgseaMultilevel (reporting NES, nominal p, adj. p across ligands, and leading-edge genes).

### Histopathology and Quantitative Immunohistochemistry

Histological examinations were performed on FFPE blocks from the middle frontal lobe. Immunohistochemistry (IHC) was performed using the following primary antibodies: anti–IBA1 (GeneTex, GTX100042, Irvine, CA, USA; 1:1000), anti–phospho-tau (clone AT8, mouse monoclonal, Thermo Fisher Scientific, MN1020, 1:1000), and anti–HLA-DR/DP/DQ (clone CR3/43, Agilent Dako, M0775, Glostrup, Denmark; 1:200). Antigen retrieval was conducted on a Dako PT Link system with low-pH solution, followed by automated staining on a Dako Autostainer Link 48 using the EnVision FLEX+ visualization system, according to the manufacturers’ instructions; all sections were counterstained with hematoxylin. Whole-slide images were acquired using a TissueScope LE120 slide scanner (Huron Digital Pathology, Waterloo, Ontario, Canada). Quantification was performed in HALO (version 2.3; Indica Labs, Albuquerque, New Mexico, USA) using the Area Quantification module. For each slide, cortex and white matter were manually annotated as ROIs; meninges, large vessels, and artefacts were excluded. For each marker (AT8, IBA1, HLA-DR/DP/DQ), the percent positive area was computed as DAB-positive area / annotated tissue area within cortex and white matter, respectively.

### Microglia Morphological Analysis

Morphometric analysis was performed on IBA1-stained whole-slide images using the Microglia Activation Module within HALO image analysis software. Gray and white matter were annotated as separate regions of interest. Within each region, microglia were detected, and features were extracted using the module’s skeletonization-based tracing, including cell count, total process length per cell (μm), process area (μm²), cell body area (μm²), and number of branch points. Detection parameters were held constant for all sections as follows: microglial cell-body diameter, 4.375–35 μm; minimum cell-body optical density (OD), 0.219; minimum process OD, 0.252; maximum process radius, 45 μm; and maximum fragmentation length, 1.06 μm. Per-cell morphometric outputs were exported and analyzed in RStudio. For distributional visualization, kernel density estimates (Gaussian kernel; default bandwidth) were used. For group comparisons, per-cell measurements were aggregated to case-level means per region (“pseudobulk” summaries), displayed as box plots, and compared between PSP and control groups using two-sided Wilcoxon rank-sum (Mann–Whitney U) tests.

### Protein-folding / ER stress assay in HMC3 microglia

HMC3 human microglial cells (ATCC, Manassas, VA, USA; CRL-3304) were cultured in DMEM, high glucose (Gibco, Thermo Fisher Scientific, Waltham, MA, USA; 11965-092) with 10% (v/v) fetal bovine serum (FBS) and 1% penicillin–streptomycin at 37 °C in a humidified 5% COl incubator. Cells were seeded at 1.0 × 10l cells per well in 12-well plates and allowed to adhere for 24 h. Tunicamycin (MedChemExpress, Monmouth Junction, NJ, USA; HY-A0098) was dissolved in DMSO, and added at final concentrations of 0.01, 0.05, 0.1, 0.5, 1, and 5 µg/mL; the control received the same final DMSO concentration (0.1% v/v). At 24 h and 48 h after treatment, the culture medium was aspirated, and cells were gently rinsed once with PBS, detached with TrypLE™ Select (Gibco, A1217701), and counted using a Countess™ II automated cell counter (Thermo Fisher Scientific). Data are displayed as mean ± SEM (n = 3 independent wells per condition). For each time point, a linear model was fitted and estimated marginal means were computed (emmeans), followed by Dunnett’s test with multivariate-t adjustment comparing each tunicamycin concentration to the DMSO control.

### LDH assay

At each time point (24 h and 48 h), prior to PBS washing, 1 µL of conditioned medium was collected from each well and immediately diluted 1:100 in LDH storage buffer (200 mM Tris-HCl, pH 7.3; 10% glycerol; 1% BSA). LDH activity in the supernatants was quantified using the LDH-Glo™ Cytotoxicity Assay (Promega, J2380, Madison, USA) according to the manufacturer’s instructions. Briefly, diluted samples were transferred to opaque white 384-well plates and mixed with LDH-Glo reagents; after incubation at room temperature, chemiluminescence (relative light units, RLU) was recorded on a FLUOstar Omega multimode plate reader (BMG LABTECH, Ortenberg, Germany) in luminescence mode (fixed gain). Each biological sample (well) was assayed in technical triplicate on the assay plate, and the mean RLU was used for analysis. In a secondary analysis, RLU were normalized to the adherent cell counts obtained from matched wells to report LDH per cell. Data are presented as mean ± SEM (n = 3 independent wells per condition). For each time point, a linear model was fitted and estimated marginal means were computed (emmeans), followed by Dunnett’s multiple comparison test with the multivariate-t adjustment, comparing each tunicamycin concentration with the DMSO control.

### Quantitative real-time PCR (qPCR)

Total RNA was extracted using the RNeasy Mini Kit (QIAGEN, 74104, Hilden, Germany) according to the manufacturer’s protocol. cDNA was synthesized with the High-Capacity cDNA Reverse Transcription Kit (Applied Biosystems, 4368814, Waltham, MA, USA). qPCR was performed using PowerUp™ SYBR® Green Master Mix (Applied Biosystems, A25776) on a LightCycler® 480 real-time PCR system (Roche Diagnostics, Basel, Switzerland). Cp values were called in the instrument software, and primer specificity was verified by melt-curve analysis (single peak). Primer sequences are provided in Source Data Extended Data Fig. 3. For Protein-folding / ER-stress experiments, cells were treated with tunicamycin (0.1 µg/mL, 48 h), with DMSO (0.1% v/v, 48h) as a control. An additional ER-stress condition with thapsigargin (MedchemExpress, HY-13433, 1 µM, 48 h) was included. Relative mRNA abundance was calculated by the 2^−ΔΔCt^, normalizing to a reference gene (GAPDH) and using the DMSO group as the calibrator(n=3 for each condition). For statistics, analyses were performed on ΔCp values. Two-group comparisons used Welch’s t test. For the three-group setting involving DMSO, tunicamycin, and thapsigargin, Welch’s t tests with Benjamini–Hochberg correction were applied within the gene/time stratum. Results are reported as mean ± SEM with n = 3 independent wells per condition.

### Cell area comparison

Cells were treated for 48 h with DMSO, tunicamycin, or thapsigargin, fixed in 4% paraformaldehyde for 15 min at room temperature. Samples were incubated with anti–HLA-DR/DP/DQ (CR3/43, Abcam, ab7856, Cambridge, UK, 1:500) in PBS with 0.05% Tween-20 at 4 °C overnight. The following day, samples were washed and incubated with AlexaFluor-488 donkey–anti-rabbit (ThermoFisher, 1:250) for 60 min and NucBlue^TM^ Fixed Cell Stain ReadyProbes^TM^ reagent (Invitrogen, P37606) for 15 minutes at room temperature. Images were obtained on a confocal microscope (ECLIPSE Ti2, Nikon, Tokyo, Japan). For each condition, three independent dishes were imaged with five non-overlapping 5 fields per dish (n = 3 dishes). In Fiji, the IBA1-positive area was measured and converted to µm² using the image scale and divided by the number of DAPI-positive nuclei in the same field to yield IBA1-positive area per cell (µm²/cell), used as a proxy for cell area. For statistics, field-level values were analyzed with a linear mixed model with treatment as a fixed effect and dish as a random intercept; post-hoc contrasts used Dunnett vs DMSO.

### Time-lapse imaging

HMC3 cells were seeded at 1.0 × 10^5^ cells per well in 12-well tissue-culture-treated plates and allowed to attach for 24 h at 37 °C, 5% COl. The medium was then replaced with DMEM/F-12 with HEPES (25 mM) (Gibco, 11330-032) supplemented with 10% FBS, 1% penicillin–streptomycin, GlutaMAX™ (Gibco, A5873601). Tunicamycin or Control (DMSO, final 0.1% v/v) was added to reach a final tunicamycin concentration of 0.1 µg/mL, and cells were incubated for a further 24 h prior to imaging. Time-lapse imaging was performed on a ZEISS Primovert inverted cell culture microscope (Carl Zeiss, Jena, Germany) equipped with a 10× objective lens. The temprature was maintained at 37 °C. A bright-field images were obrained 5MP USB 2.0 Digital Eyepiece Camera (OMAX, A3550S) controlled by ToupView were used. Single-plane images were acquired at 1-min intervals for 12 h (from 24 to 36 h post-treatment).

## Notes

### Competing Interest Statement

The authors have declared no competing interest.

