## Supplementary material for "Protein folding stress shapes microglial phenotype in progressive supranuclear palsy": Figure S1-4

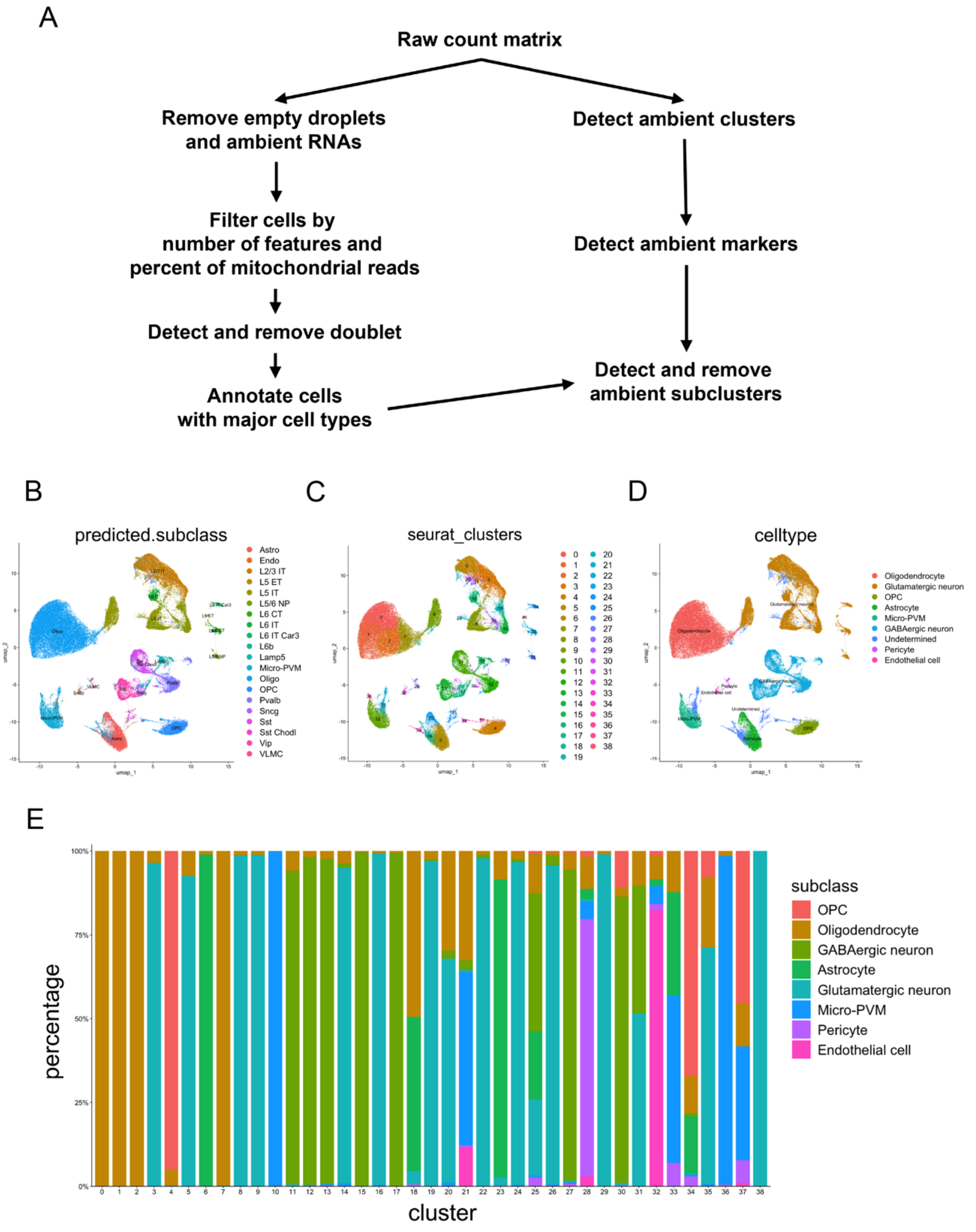


**Figure S1 Quality Control and Cell annotation**

(A) Workflow of snRNA-seq quality control. (B-D) UMAP visualizations of snRNA-seq nuclei after CellBender-based ambient RNA removal, filtering by nFeature_RNA and percent.mt, and doublet removal. (B) Reference-mapped subclasses (predicted.subclass); (C) Seurat clusters; (D) manually curated major cell types (oligodendrocyte, glutamatergic neuron, oligodendrocyte progenitor cell (OPC), astrocyte, microglia/perivascular macrophage, GABAergic neuron, pericyte, endothelial cell, undetermined). (E) Proportion of major cell types in each Seurat cluster.

### Figure S2

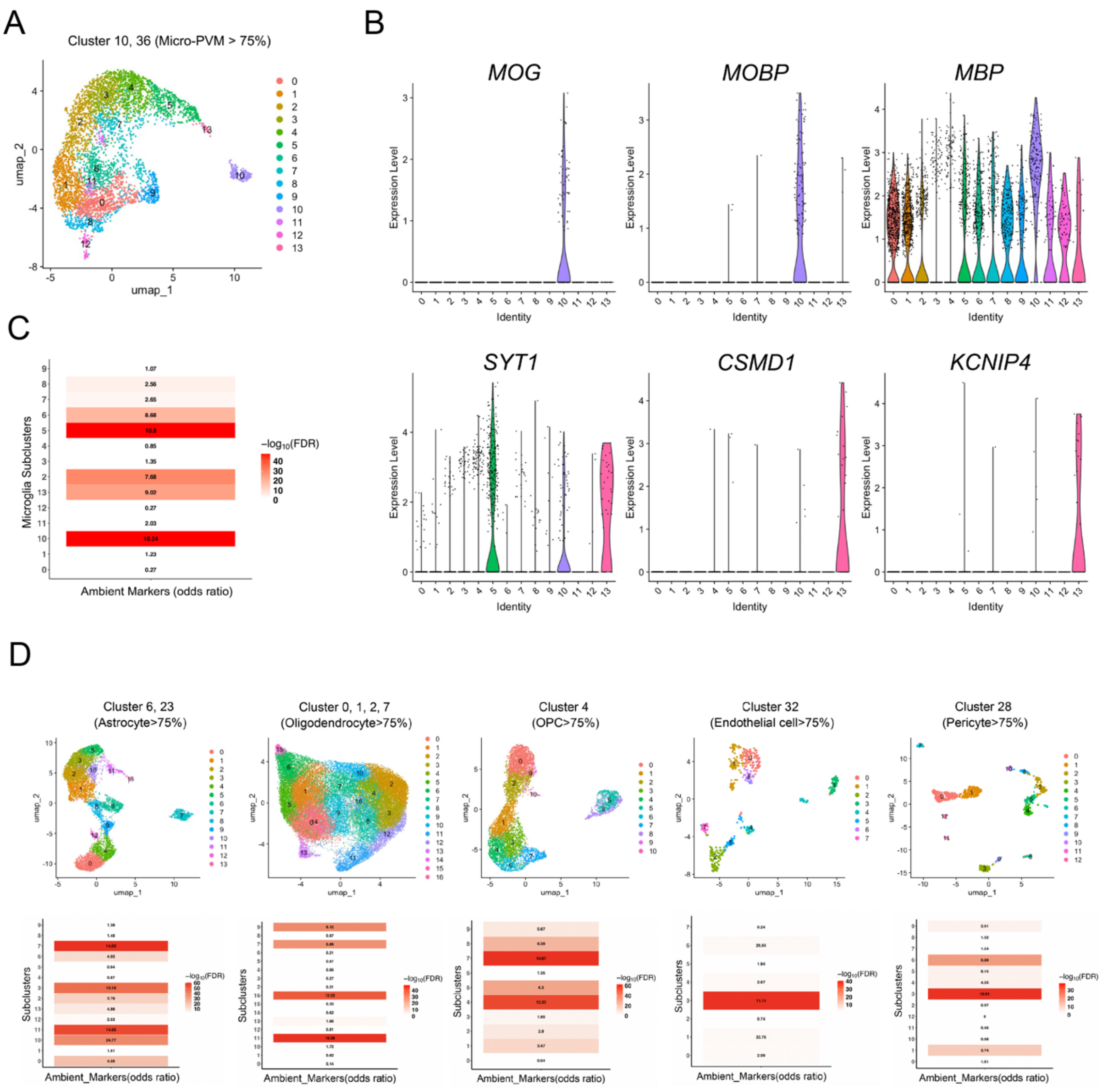


**Figure S2 Ambient Cluster Cleaning**

(A) UMAP of the microglia/perivascular macrophage subset. Nuclei from Seurat clusters 10 and 36, in which Micro-PVM accounted for >75% of cells, were extracted and reclustered; colors indicate subclusters (0–13). (B) Violin plots showing expression of canonical oligodendrocyte markers (*MOG*, *MOBP* and *MBP*) and neuronal markers (*SYT1*, *CSMD1* and *KCNIP4*) across microglia/perivascular macrophage subclusters (0–13). (C) Enrichment of ambient marker genes across microglial subclusters. Horizontal bars show the odds ratio for enrichment of ambient markers in each subcluster; color indicates $-\log_{10}$(adjusted p). Subclusters with adj. $p < 0.001$ and odds ratio > 3 were classified as ambient and excluded from downstream analyses. (D) Cell-type–specific subclustering and ambient cluster removal.

Figure S3

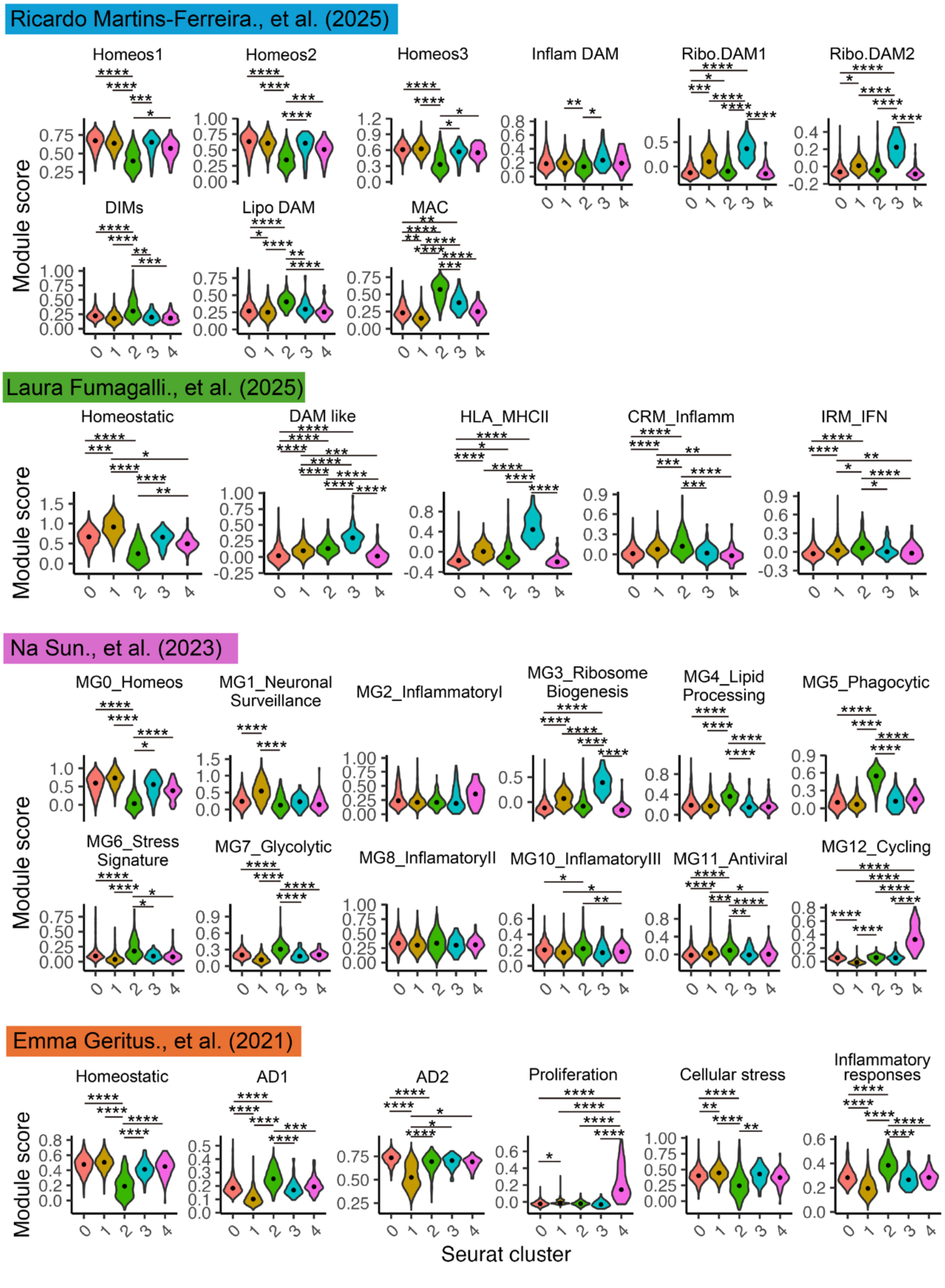


**Figure S3 Module scores for previously reported microglial gene sets**

Violin plots showing module scores for 32 literature-derived microglial gene sets across Microglia-PVM subclusters (SC0–SC4). Per-cell scores were computed with Seurat AddModuleScore and then aggregated to donor × subcluster means. Cluster effects were tested with a weighted linear mixed model including cluster as a fixed effect and donor as a random intercept; pairwise contrasts between subclusters were obtained with emmeans and Benjamini–Hochberg correction within each module. ****adj. $p < 0.0001$, ***adj. $p < 0.001$, **adj. $p < 0.01$, *adj. $p < 0.05$.

Figure S4

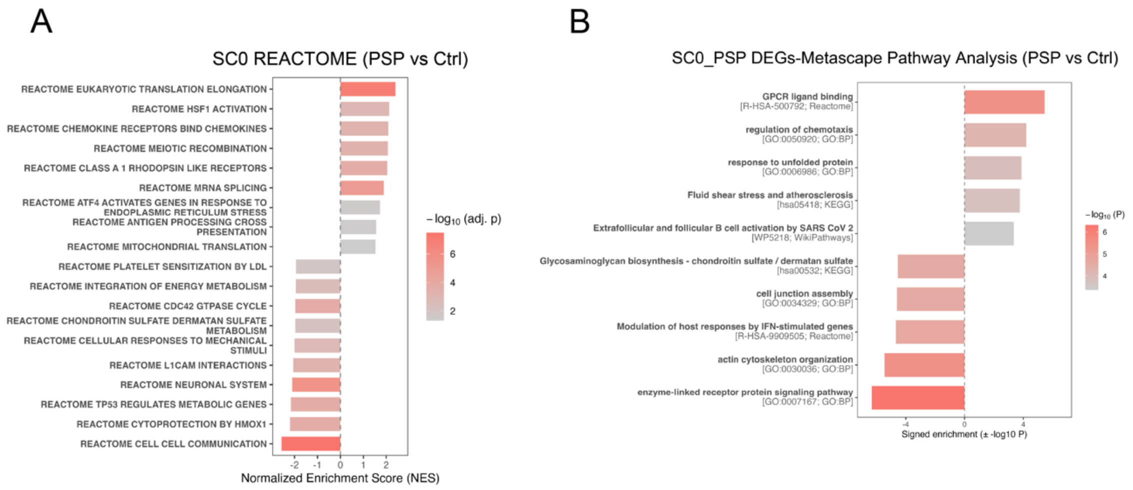


**Figure S4 Extended pathway analyses of SC0 microglia (PSP vs Ctrl)**

(A) Gene-set enrichment analysis using Reactome pathways for SC0 microglia (PSP vs Ctrl). Bars show the normalized enrichment score (NES) for significantly enriched pathways, with color indicating $-\log_{10}$(adjusted p). Positive NES indicates relative upregulation in PSP. (B) Over-representation analysis of SC0_PSP differentially expressed genes (PSP vs Ctrl) performed with Metascape; color encodes $-\log_{10}$(p value).

**Video S1 Effect of tunicamycin on HMC3 microglial motility**

Time-lapse imaging of HMC3 cells treated with vehicle control (DMSO, right) or tunicamycin (0.1 μg/mL, left). Images were acquired at 1-min intervals from 24 to 36 h post-treatment.
